# Virtual clinical trial reveals significant clinical potential of targeting tumour-associated macrophages and microglia to treat glioblastoma

**DOI:** 10.1101/2024.12.06.627263

**Authors:** Blanche Mongeon, Morgan Craig

## Abstract

Glioblastoma is the most aggressive primary brain tumour, with a median survival of just fifteen months with treatment. Standard-of-care (SOC) for glioblastoma consists of resection followed by radio- and chemotherapy. Clinical trials involving PD-1 inhibition with nivolumab in combination with SOC failed to increase overall survival. A quantitative understanding of the interactions between the tumour and its immune environment driving treatment outcomes is currently lacking. As such, we developed a mathematical model of tumour growth that considers cytotoxic CD8+ T cells, pro- and antitumoral tumour-associated macrophages and microglia (TAMs), SOC, and nivolumab. Our results show that PD-1 inhibition fails due to a lack of CD8+ T cell recruitment during treatment explained by TAM-driven immunosuppressive mechanisms. Using our model, we studied five TAM-targeting strategies currently under investigation for solid tumours. Our model predicts that while reducing TAM numbers does not improve prognosis, altering their functions to counter their protumoral properties has the potential to considerably reduce post-treatment tumour burden. In particular, restoring antitumoral TAM phagocytic activity through anti-CD47 treatment in combination with SOC was predicted to nearly eradicate the tumour. By studying time-varying efficacy with the same half-life as the anti-CD47 antibody Hu5F9-G4, our model predicts that repeated dosing of anti-CD47 provides sustained control of tumour growth. Thus, we propose that targeting TAMs by enhancing their antitumoral properties is a highly promising avenue to treat glioblastoma and warrants future clinical development. Together, our results provide proof-of-concept that mechanistic mathematical modelling can uncover the mechanisms driving treatment outcomes and explore the potential of novel treatment strategies for hard-to-treat tumours like glioblastoma.

## Introduction

Glioblastoma is an aggressive brain tumour that accounts for about 46.1% of all malignant primary brain tumours^1^. Standard-of-care (SOC) for glioblastoma consists of maximal surgical resection followed by radiotherapy and temozolomide (TMZ) chemotherapy, resulting in a median overall survival after SOC of just barely fifteen months^2^. There is considerable interest in developing immunotherapies against glioblastoma^3^. It was thought that immune checkpoint inhibitors (ICIs) targeting the immunosuppressive PD-1/PD-L1 signaling axis would be successful, given that a large proportion of newly diagnosed adult glioblastomas express PD-L1^4^. However none of the multiple clinical trials of nivolumab, an anti-PD-1 monoclonal antibody, have resulted in improvements to overall survival^5–7^.

Recent studies highlight that glioblastoma immune suppression limits ICI efficacy^8^. Tumour-associated macrophages and microglia (TAMs) are often the most abundant immune cells in the tumour microenvironment (TME)^9^. As TAMs kill tumour cells *in vitro*^10^, they were initially believed to be solely antitumoral. There is however growing evidence that some TAM subpopulations promote tumour progression^11^ and interfere with antitumoral therapy^9^. Indeed, an increased abundance of TAMs in the TME is associated with a decrease in patient survival in many cancers^9^. TAMs exist on continuum of polarization states with two extremes (M1 and M2)^12^. M1-like TAMs have greater antitumoral functions, including tumour cell phagocytosis^13^ and high antigen presenting capacity^14^, while M2-like TAMs promote an immunosuppressive TME through the secretion of cytokines that reduce effector T cell functions and induce immune checkpoint expression^12^. In glioblastoma, high levels of immunomodulatory factors like transforming growth factor beta (TGF-β)^15^ and interleukin-10 (IL-10)^16^ contribute to immunosuppressive TAM phenotypes^8^. Further, patients in the Cancer Genome Atlas (TCGA) with a higher expression of M2-like TAM signatures were found to have significant reductions in predicted survival^17^. Thus, an M2-like phenotypic skew results in low adaptive immune cell recruitment and limits the efficacy of immune checkpoint blockade^18^. As such, TAM-mediated immunosuppression is a potential target to improve SOC outcomes and design effective ICI regimens for glioblastoma^8^.

As highlighted by the nivolumab clinical trial failures and the mere 5% success rate from first-in-human trials to registration^19^, the development of new drugs in oncology is difficult and costly. Approaches enabling the early identification of strategies with the potential to succeed in trials are highly needed. Mathematical modelling and virtual clinical trials (VCTs) allow us to study novel therapeutic approaches in a robust and cost-effective way^20^. Zhang et al.^21^ studied the evolution of resistance in metastatic castrate-resistant prostate cancer treated with abiraterone using an evolutionary game theory model. Their model predicted that stopping abiraterone treatment when prostate-specific antigen (PSA) levels fall below 50% of the original value and withholding the drug until PSA returns to its original levels would suppress proliferation of androgen-independent cancer cells. This strategy was then validated in a pilot clinical trial where median time to progression increased from 16.5 to 27 months, and cumulative drug use reduced by 47% compared to standard dosing^21^.

Given the relevance of TAMs to ICI success in glioblastoma, we developed a mathematical model of glioblastoma and its TME to study treatments combining SOC, nivolumab and various TAM-targeting agents. Our results show that altering TAM protumoral functionalities is a more promising therapeutic avenue than reducing their number in the TME. Encouragingly, enhancing the phagocytic activity of M1 TAMs was predicted to significantly increase survival in a virtual patient cohort. This work therefore supports the ongoing development of TAM-targeting agents for their potential to deliver needed benefits for glioblastoma treatment.

## Materials and Methods

### Mathematical model of glioblastoma

To investigate glioblastoma treatment, we developed a modular mathematical model describing tumour growth, innate and adaptive immune responses, and surgical resection, radiotherapy, chemotherapy, ICIs, and TAM-targeting agents (**Figure 1**). Complete model equations are provided in the Supplementary Information (SI), with state variables and units listed in **Table 1**.

**Table 1.**
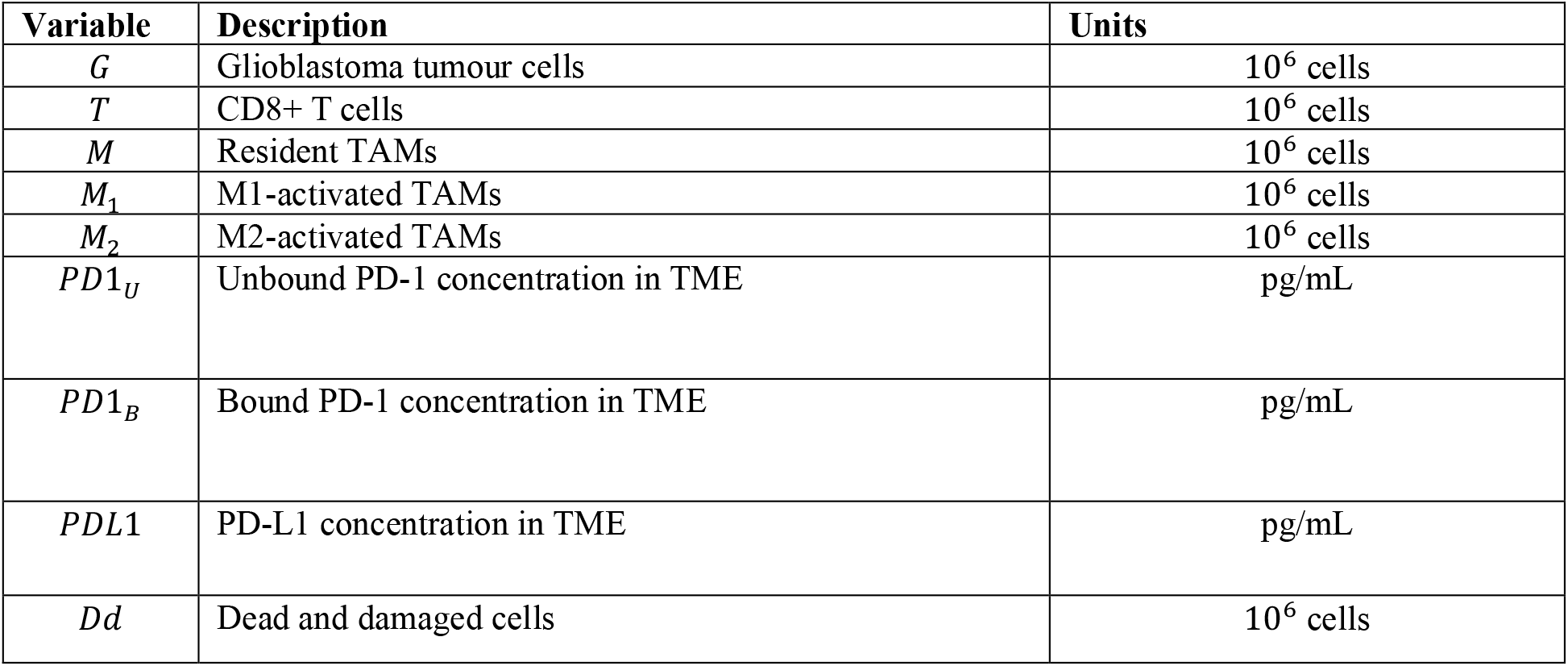
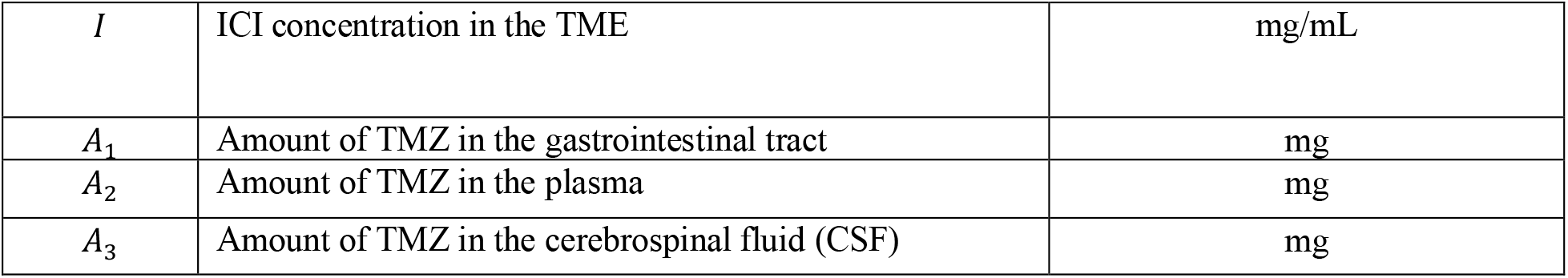
Variables of the mathematical model, with their description and units.

**Figure 1.**
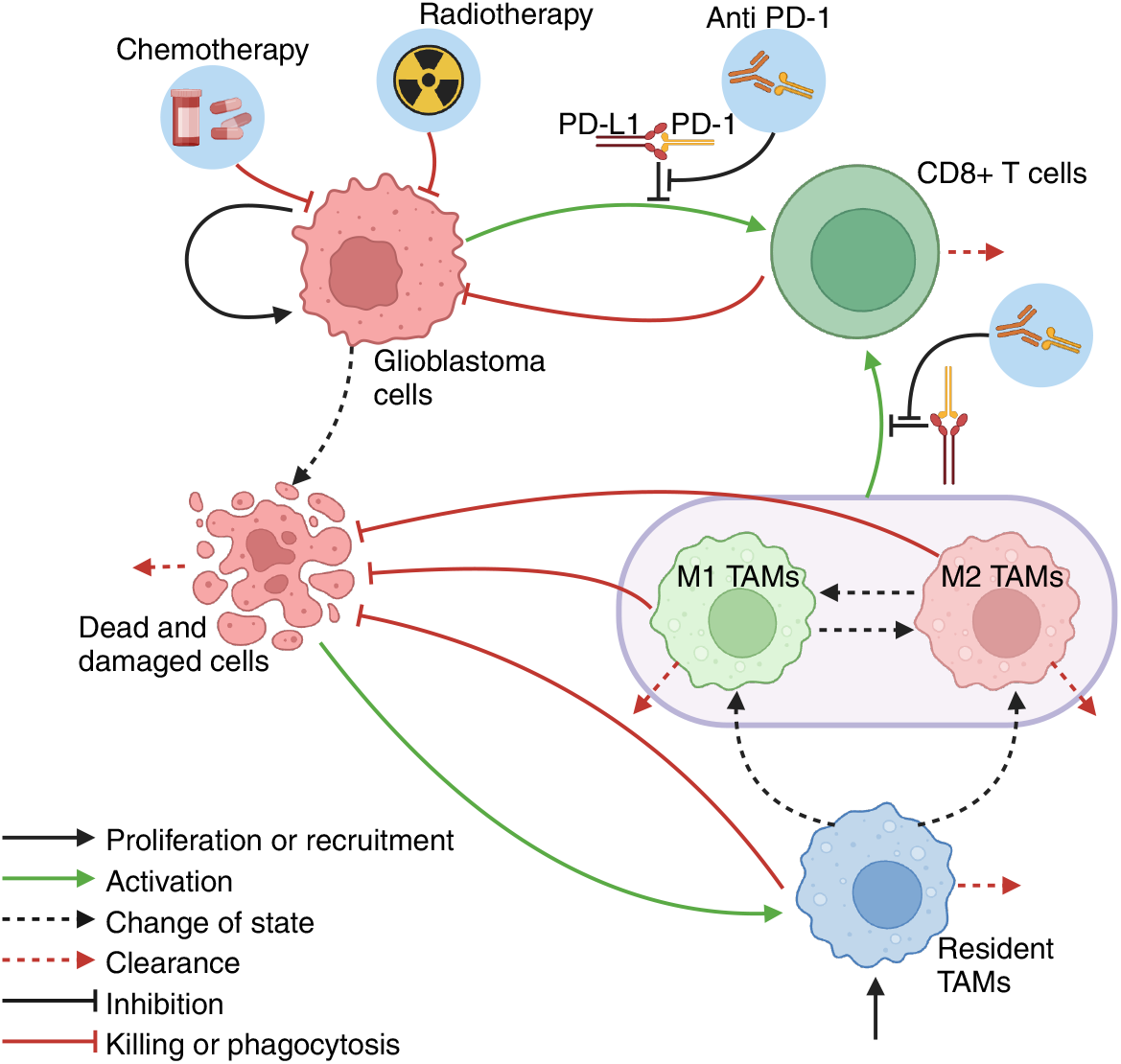
Mathematical model of glioblastoma growth, TME, and treatment. Glioblastoma cells proliferate and are killed by cytotoxic CD8+ T cells. Resident tumour-associated macrophages and microglia (TAMs) are recruited to the TME at a constant rate, and activated by dead and damaged tumour cells and activated M1 and M2 TAMs. M1 and M2 TAMs have anti- and protumoral functions, respectively. All TAMs can phagocytose dead and damaged cells. CD8+ T cells are activated by glioblastoma cells and TAMs, depending on the M2:M1 ratio. This activation is inhibited by PD-1/PD-L1 signalling, mimicking the exhaustion of CD8+ T cells. Chemotherapy and radiotherapy act directly on tumour cells, while ICIs prevent binding between PD-1 receptors on CD8+ T cells and PD-L1 expressed on tumour cells and TAMs. Made with BioRender.

We assumed Gompertzian tumour growth dynamics, with untreated tumour cells growing biexponentially to their carrying capacity. Further, we modelled a constant source of resident TAMs activated through phagocytic activity or direct interaction with activated TAMs. For simplicity, we considered only dual TAM polarization so that resident TAMs are either M1 or M2 once activated. The parameter *q* accounts for the fraction of resident TAMs activated with the M2 phenotype and hence the M2 TAM bias. We modelled phenotypic switching of activated TAMs^11,14^ and natural death to occur at constant rates.

Tumour cells can be killed by cytotoxic CD8+ T cells. Primed CD8+ T cells die through natural death and are recruited to the tumour site by glioblastoma cells and activated TAMs. Assuming that an over-abundance of tumour cells restricts movement within the tumour architecture and saturates the immune response, we modelled saturable recruitment using Michaelis-Menten kinetics^22^. Since M1 and M2 TAMs have anti- and protumoral properties, respectively, we considered the activation of CD8+ T cells to depend on the ratio of M2 to M1 TAMs^12^. Thus, when M2 TAMs far outnumber M1 TAMs, there is no CD8+ T cell recruitment, whereas activated TAMs recruit CD8+ T cells if not.

Unbound PD-1 receptors (*PD*1_*U*_) are free to bind to PD-L1 receptors to inhibit CD8+ T cell cytotoxicity. Dynamics of bound and unbound receptors (see **Treatment with an ICI**) were modelled by tracking the concentration of PD-1 receptors on new CD8+ T cells in the TME, with PD-1 receptors removed as CD8+ T cells die^22^. We considered glioblastoma cells^4^ and all three types of TAMs^9^ to express PD-L1 receptors and modelled their dynamics similar to PD-1 receptors. As in previous work^22,23^, T cell exhaustion due to PD-1/PD-L1 complexes was modelled as suppression of T cell activation:

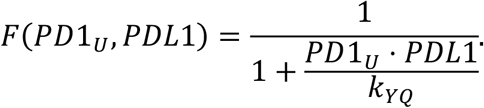

Here, more PD-1/PD-L1 complexes are formed as *PD*1_*U*_ and *PDL*1 concentrations increase, with *F*(*PD*1_*U*_, *PDL*1) saturating to 0.

Finally, we tracked the effects of CD8+ T cell cytotoxicity and TAM phagocytosis by explicitly accounting for damaged and dead cells resulting from CD8+ T cell killing.

### Treatment with SOC

SOC consists of surgical resection followed by concomitant 42-day cycle with a five-days-a-week 2 Gy/day radiation fractions and daily oral TMZ (75 mg/kg). After a four-week break, patients receive six adjuvant 28-day cycles of chemotherapy with TMZ given on days one to five in doses ranging from 150 mg/kg to 200 mg/kg (**Figure 2A**). Based on clinical measurements, we modelled resection as a partial reduction to all cell populations (see **SI** section **Treatment with resection**). We also used a direct tumour volume reduction^24^ to model radiotherapy based on the linear-quadratic dose response model describing cell survival probability after a radiation fraction^25^. We integrated a three-compartment population pharmacokinetic (PopPK) model^26^ (**Supplementary Figure 1A**) to track TMZ concentrations in the gastrointestinal tract, plasma, and cerebrospinal fluid (CSF) over time (see **SI** section **Treatment with temozolomide**), with the CSF as site of action. Although TMZ is considered to be primarily cytostatic, we assumed it induces cell death in proliferating cells by arresting their cell cycle^27^ and modelled TMZ as a cytotoxic drug (see **SI** section **Treatment with temozolomide**).

**Figure 2.**
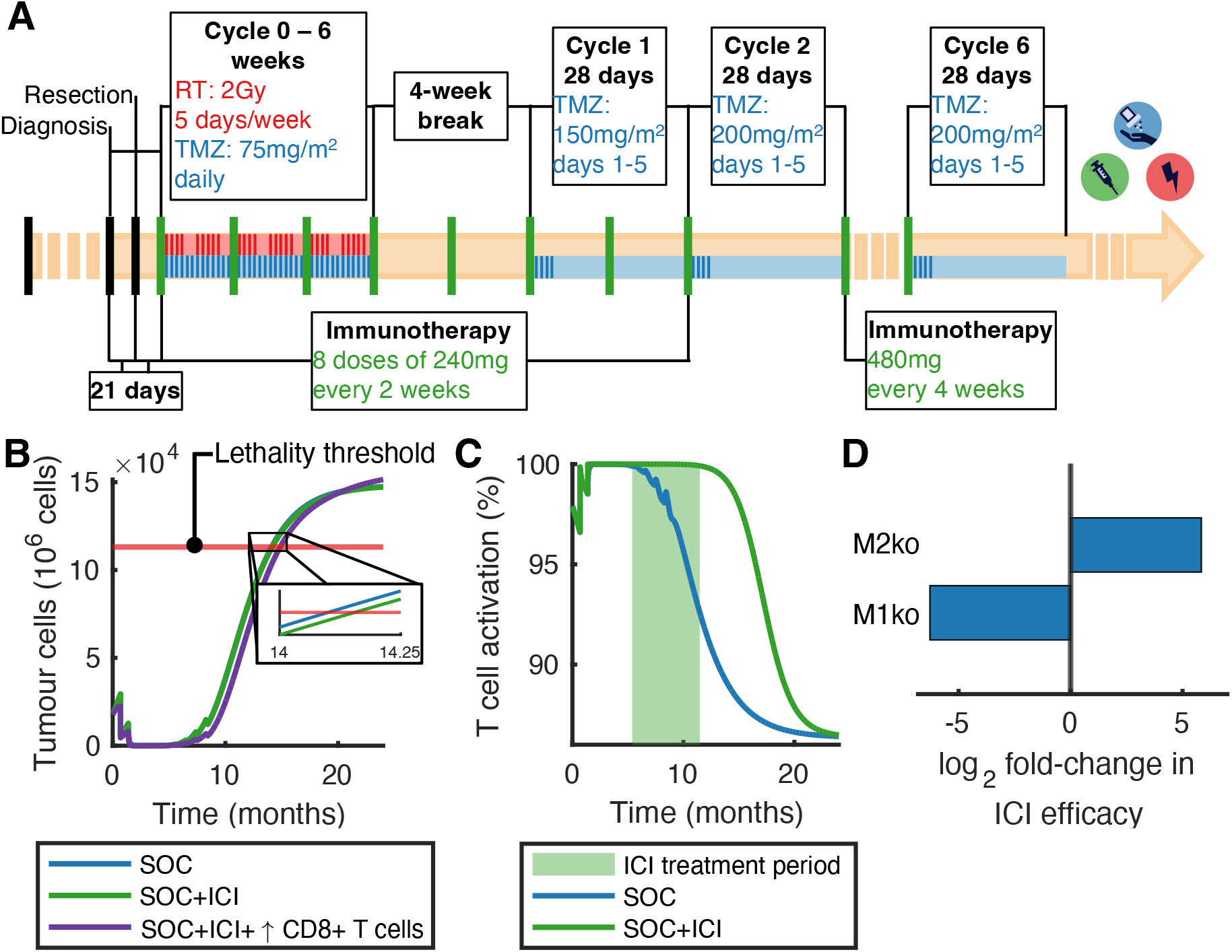
Lack of CD8+ T cell recruitment to the TME drives ICI clinical trial failures. **A)** Simulated treatment schedules. Resection was simulated 21 days after diagnosis with chemoradiotherapy cycles starting 3 weeks after surgery. TMZ was administered orally daily while radiotherapy given weekly on days 1 to 5, for a total of 6 weeks. After a four-week break, oral TMZ was administered in increasing doses on days 1 to 5 for six 28-days cycles of chemotherapy. For ICIs, we considered eight doses of 240 mg given at two-week intervals from the start of the treatment, with 480 mg doses given every four weeks to the end of chemotherapy. **B)** Tumour growth after treatment with SOC (blue) versus SOC with an ICI (green). Red line: cell count at which the tumour is considered lethal. The addition of the ICI resulted in a survival time gain of about 2 days. However, increasing CD8+ T cell recruitment (purple) resulted in increased survival. **C)** CD8+ T cell activation during SOC (blue) and SOC+ICI (green) as described by 1 − *F*(*PD*1_*U*_, *PDL*1), see section Mathematical model of glioblastoma. Shaded green region shows the treatment period during which we simulate nivolumab administration. Anti-PD-1 ICI prevents the suppression of CD8+ T cell activation by PD-1/PD-L1 binding. **B-C)** Time refers to time after diagnosis. **D)** M2 knockout during SOC+ICI treatment resulted in increased nivolumab efficacy (i.e., *S*(*SOC* + *ICI*)) while M1 knockout caused decreased efficacy.

### Treatment with immune checkpoint inhibitors

For safety, nivolumab is approved as a flat dosage regimen of either 240 mg administered every two weeks or 480 mg every four weeks^28^. We modelled nivolumab administration as a one-hour intravenous infusion (see **SI** section **Immune checkpoint inhibitor dosing**). As in the phase III clinical trial by Lim et al.^29^, we considered an ICI dosing regimen of 240 mg every two weeks for eight cycles starting at the first dose of TMZ, followed by 480 mg administrated every four weeks until the end of chemotherapy (**Figure 2A**).

### Potential strategies targeting tumour-associated macrophages and microglia

Without treatment, glioblastoma cells escape TAM phagocytosis through the expression of “*don’t eat me*” signals such as Cluster of Differentiation 47 (CD47)^9,30^. Targeting TAMs can take the form of reducing their numbers in the TME and/or altering their function. To treat solid tumours, there are four main strategies: 1) delete TAMs, 2) inhibit TAM recruitment to the TME, 3) restore TAM phagocytotic activity, and 4) reprogram protumoral TAMs into antitumoral immune cells^9^. To model these, we introduced five parameters describing the efficacy of TAM-targeting molecules, as described below (see **SI** section **Targeting tumour-associated macrophages and microglia** for complete description).

Colony-stimulating factor-1 receptor (CSF1) binds to its receptor (CSF1R) to regulate TAM survival, proliferation, and differentiation^31^. CSF1R blockade via an anti-CSF1R mAB such as Emactuzumab^32^ could thus deplete TAMs^9^, which we modelled by increasing the rate of TAM apoptosis. Chemokine (C-C motif) ligand 2 (CCL2) is produced by various central nervous system resident cells^33^ and is a chemoattractant for monocytes and other cells^9^. TAM recruitment to the TME could thus be inhibited by targeting CCL2 and its receptor (CCR2)^9,33^. Systemic injection of the anti-CCL2 mAB Carlumab^9^ in prostate cancer mouse models showed reduced infiltration of CD68+ macrophages^34^. We modelled this by inhibiting TAM recruitment to the TME through decreasing the constant source of TAMs. CD47 has been found to be a highly expressed “*don’t eat me*” signal on glioblastoma cells^30^. We modelled the effect of an anti-CD47 mAB such as Hu5F9-G4^35,36^ by enabling increased phagocytosis of tumour cells by M1 TAMs (with no effect on protumoral M2 TAMs). Finally, reprogramming the TME to be antitumoral could be achieved by using Toll-like receptors (TLR) agonists like imiquimob^9^ and 852A^37^ to polarize TAMs towards a pro-inflammatory phenotype^9^. To model TAM reprogramming with TLR agonists, we increased the switching rate from M2 to M1 and decreased (increased) the TME bias towards M2 (M1) activated TAMs.

### Parameter estimation and initial conditions

Model parameters were estimated from the literature or fit to publicly available data. Full details are included in the **Supplementary Information**, with parameter values listed in **Supplementary Tables 2** to **9**.

Parameters in the linear quadratic radiotherapy relationship were taken from Franken and al.^38^ based on VU-109 glioma cell data after exposure to TMZ-radiation treatment. The TMZ sigmoidal effects curve was fit to data from cell viability assays performed by Lo Dico and al.^39^ using *lsqnonlin* in MatlabR2024a^40^. The ICI model was parametrized according to Storey et al.^22^ and rescaled to our units. The tumour intrinsic growth rate and the maximal killing rate of glioblastoma cells by CD8+ T cells were fit to survival time data reported by Stupp et al.^2^ using *lsqnonlin* in in MatlabR2024^40^ (see **SI** section **Detailed parameter estimation calculations**).

To model tumour initiation, we assumed an initial tumour volume of 5 cubic millimeters (i.e., *G*(0) = 5 · 10^6^ tumour cells). Initial cell-to-cell ratios were obtained from Surendran et al.^18,^ and Karimi et al.^41^ based on imaging mass cytometry analyses of glioblastoma biopsies and resections (see **SI** section **Initial conditions**).

### Predicting treatment outcomes and efficacy analysis

Using our parametrized model, we studied the treatment strategies described above. We considered post-treatment time to be 7 days after the last administration of a TMZ dose. The efficacy of a novel therapy 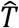 was calculated as the percent change of the post-treatment tumour cell count compared to SOC:

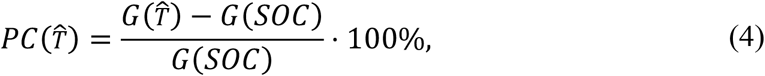

where G(T^∗^) is the tumour cell count after treatment *T*^∗^.

Supplementary efficacy conferred by 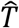, denoted by 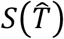, was defined as the percent decrease in tumour burden compared to SOC, i.e., 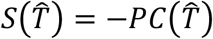. Hence, if a treatment 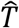 was more effective than SOC, 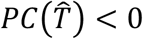 and 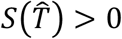, and vice versa. Note that 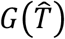 must be greater or equal to 0 and that 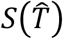 is upper bounded by 100%, representing complete eradication of the tumour.

### Parameter sensitivity analysis

To identify the parameters most impacting predicted outcomes, we performed a global sensitivity analysis using Extended Fourier Amplitude Sensitivity Test (eFAST). We evaluated the sensitivity of our model to intrinsic tumour growth, the activity of the adaptive and innate immune systems, and ICI binding efficacy. To reduce the dimensionality of the analyzed parameter set, we assumed that if our model was sensitive to the maximal rate, then it was also sensitive to its corresponding interaction and only varied the maximal rate and not the half-effect concentration in interaction terms in Michaelis-Menten relationships. We measured changes to model outputs by considering the tumour burden after SOC and the supplementary efficacy 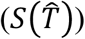 of a treatment combining SOC and nivolumab (i.e., *S*(SOC + ICI)). To identify significantly sensitive parameters, we performed a two-sample t-test using a dummy variable not in our model^42^. We defined sensitive parameters as those with a total-order sensitivity index higher than that of the dummy parameter, and resampled total-order sensitivity index distributions significantly different from that of the dummy parameter. Statistical significance was determined at α = 1%^42^.

### Generating a virtual cohort of patients and running a virtual clinical trial

We generated a cohort of virtual patients by varying tumour-specific characteristics based on the results of our global sensitivity analysis. Each virtual patient is a set of parameters fixed or sampled from predefined values. We defined log-normal distributions for tumour and TME specific parameters and integrated log-normal distributions for the TMZ pharmacokinetics per the PopPK model^26^ (see **SI**). For our VCT, we generated 900 virtual patients. If the parameter set resulted in a tumour reaching the size at diagnosis within ten years, we included it in our virtual cohort as a plausible parameter set; if not, we discarded it and resampled.

### Statistical analysis

A two-sided Wilcoxon rank sum test using *ranksum* in MatlabR2024a^40^ was used to statistically compare two distributions. Statistically significance was defined at the α = 1% level.

To identify patient-specific parameters with the highest impact on treatment efficacy, we normalized each patient’s supplementary efficacy 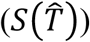 by their untreated tumour growth over the same period. This allowed us to evaluate the impact of the parameter independent of the aggressivity of tumour growth. We then segregated the virtual patient cohort into quartiles based on their normalized *S* values and defined the worst and best responders as the patients in the first and third quartiles, respectively. To identify parameters distinguishing best from worst responders, we also used an effect size measure by calculating the overlapping index 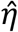^43^. This index is a similarity measure that evaluates the overlap between two distributions (η = 0 represents two completely separate distributions versus identical distributions when *η* = 1). We defined parameters distinguishing best from worst responders if the two-sided Wilcoxon rank sum test was statistically significant with 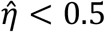.

Statistically significant differences between two Kaplan-Meier survival curves were evaluated using a log-rank test using the function *MatSurv*^44^ in MatlabR2024a^40^.

## Results

### Intrinsic immunosuppression drives glioblastoma ICI clinical trial failures

To investigate mechanisms of treatment failure, we first simulated SOC + ICI trials with surgical resection occurring 21 days post-diagnosis and chemoradiotherapy beginning three weeks later^2^ (**Figure 2A**).

Under SOC with no ICI, our model predicted a typical glioblastoma survival time of 14.10 months after diagnosis (see **SI** section **Predicting survival time**), within the 95% confidence interval (13.2 to 16.8 months) of the median survival time reported by Stupp et al.^2^. Our model further predicted a negligible survival time increase of 2 days for SOC in combination with nivolumab (**Figure 2B**), consistent with clinical trials of ICI treatment in combination with chemoradiotherapy^5,29^.

To understand why nivolumab does not increase overall survival relative to SOC, we focused on CD8+ T cells within the TME. We found that anti-PD-1 restores cytotoxic CD8+ T cell activation (**Figure 2C**), but too little to cause a significant increase in survival time due to a lack of CD8+ T cell recruitment. To validate this hypothesis, we increased the number of CD8+ T cells in the TME by simulating a constant source ten times higher than at homeostasis and found longer survival times due to increased cytotoxicity (**Figure 2B**). This is coherent with our parameter sensitivity analysis which showed that the supplementary efficacy of ICIs was not sensitive to its blocking efficacy (**Supplementary Figure 5B**). These findings suggest that ICI clinical trial failures in glioblastoma are due to immunosuppressive mechanisms that prevent CD8+ T cell recruitment.

These results motivated us to study immune escape through TAM polarization and its impact on survival and efficacy. For this, we simulated SOC+ICI in a tumour with a TME depleted of M1 and M2 TAMs (see **SI** subsection **Tumour growth in M1- and M2-knockout models**). Compared to average simulations, knockout of M1 TAMs during treatment was predicted to increase nivolumab supplementary efficacy (i.e., *S*(SOC + ICI)), which decreased during M2 knockout (**Figure 2D**), resulting in a survival time difference of approximately one month (M1 knockout: 14.06 months, M2 knock out: 14.99 months). Interestingly, the median glioblastoma survival time more closely resembled that of an M2 TAM-depleted tumour, which is consistent with an average glioblastoma having a high M2:M1 ratio of around 3.2^45^. When M2 TAMs were removed entirely at tumour initiation, tumours were predicted to stabilize below the lethal threshold (**Supplementary Figure 3**). This led us to hypothesize that TAM polarization and functionality impact ICI efficacy and success, and ultimately overall survival.

### Enhancing phagocytosis is a promising immunotherapy against glioblastoma

Given their protumoral role, we sought to predict the potential of TAM-targeting strategies to treat glioblastoma in combination with SOC with or without nivolumab (see **Methods** section **Targeting tumour-associated macrophages and microglia**). Counterintuitively, our model predicted that reducing the number of TAMs in the TME by increasing their death rate/removal could induce a small increase in tumour size after treatment (**Supplementary Figure 6A**). This is likely due to a slight increase in the M2:M1 ratio, which reduces adaptive immune activation by innate cells (**Figure 3D**). Indeed, targeting M2 TAMs by only increasing their death rate resulted in a tumour cell decrease of over 12% (**Supplementary Figure 6C**). This underlines the significant role the ratio of protumoral to antitumoral TAMs in treatment efficacy. Further, it explains why inhibiting the recruitment of resident TAMs to reduce the number of protumoral TAMs was also found to be ineffective (**Supplementary Figure 6B**), as this strategy reduces the number of M1 and M2 TAMs without altering the M2:M1 ratio driving the activation of CD8+ T cells by TAMs (**Figure 3D**).

**Figure 3.**
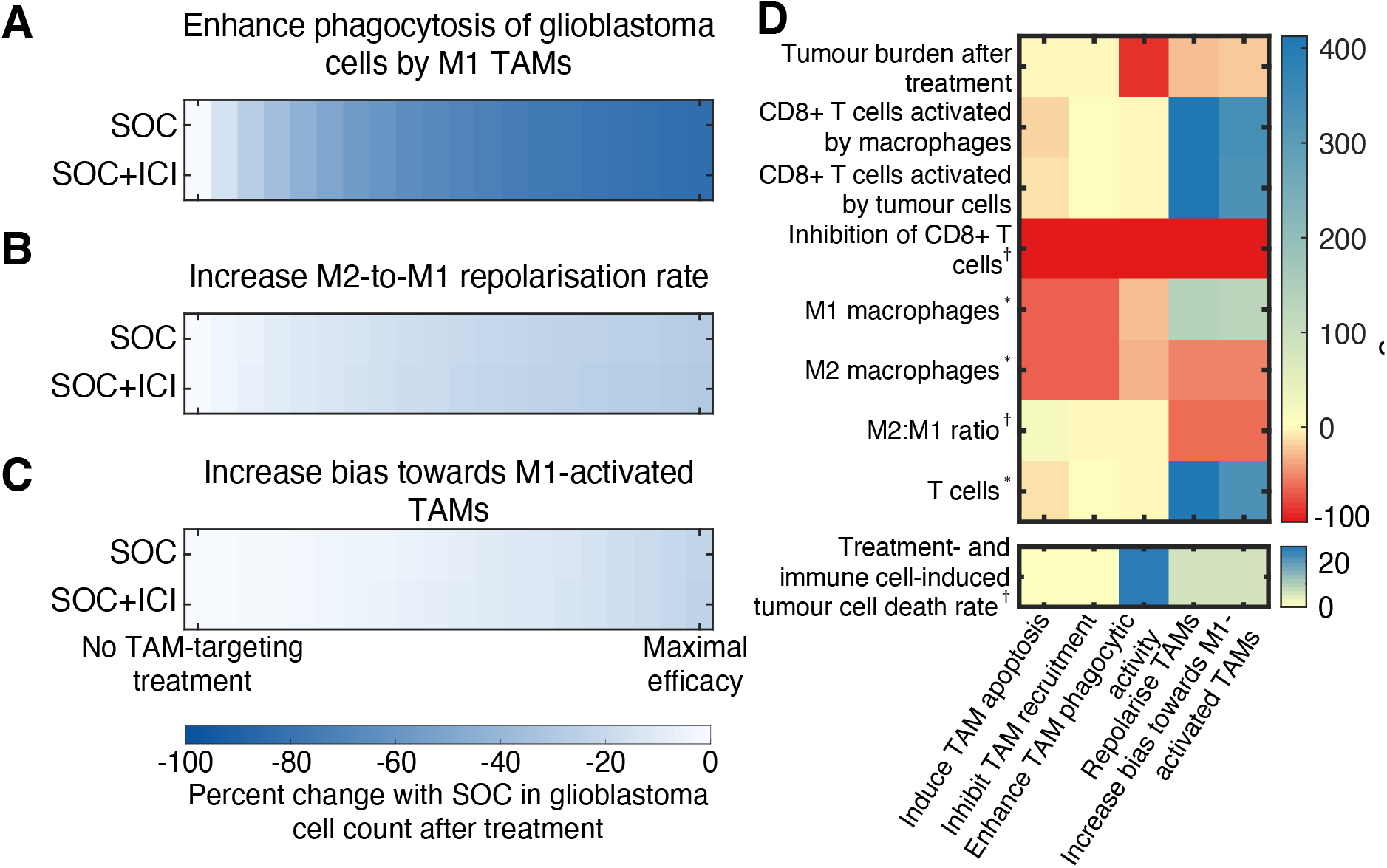
Targeting tumour-associated macrophages to alter their functional properties improves treatment outcomes. **A-C)** Efficacy of TAM-targeting treatments in combination with SOC (top rows) and SOC+ICI (bottom rows) relative to SOC. We varied the efficacy of the TAM-targeting agent between no effect and maximal (see Methods). Heatmaps report the percent change in tumour cell count one week after treatment compared to SOC. Protumoral TAM functions can be altered by A) increasing the bias towards M1-activated TAMs, B) increasing the repolarization rate of M2 TAMs into M1 TAMs, or C) enhancing the phagocytic activity of glioblastoma cells by M1 TAMs. **D)** Heatmap showing the percent change in different metrics with SOC for five TAM-targeting strategies (x-axis) in combination with SOC and ICI. The killing rate of tumour cells was defined as the fraction of glioblastoma cells removed by resection or killed by radiotherapy, chemotherapy, CD8+ T cells or phagocytosis. Decreasing the number of TAMs in the TME by either inducing their apoptosis or inhibiting their recruitment had no impact on the M2:M1 ratio and, thus, on tumour burden after treatment. Decreasing the M2:M1 ratio by increasing M2-to-M1 repolarisation rate or the TME bias towards M1-activated TAMs led to an increase in CD8+ T cells, translating to a small increase in the average killing rate of glioblastoma cells. The increase in the average rate of treatment- and immune cell-induced tumour cell death, resulting in a decrease in post-treatment tumour burden, is larger when the phagocytic activity of M1 TAMs is enhanced, as this strategy has a direct effect in tumour cells. ∗: area under the curve of the cell population during treatment, †: average value during the treatment period.

We next wanted to understand whether altering the functional properties of TAMs instead of their numbers could improve therapeutic outcomes. We found that increasing the M1 bias (i.e., inducing an antitumoral phenotype to newly activated TAMs) resulted in a 22.53% decrease of the maximal final tumour size without nivolumab and 23.35% decrease with anti-PD-1 (**Figure 3C**). Though small, this 0.82% difference in the maximal efficacy when nivolumab is added to the combination was over ten-times the difference between SOC and SOC+ICI (0.06%). Similarly, increasing the M2-to-M1 repolarization rate resulted in maximal decreases of 27.27% (without nivolumab) and 28.25% (with nivolumab) to the final tumour size (**Figure 3B**).

Crucially, our results suggest that enhancing tumour cells phagocytosis by M1 TAMs is significantly more effective versus SOC, resulting in a maximal decrease in an 82.47% reduction of the post-treatment glioblastoma cell count without nivolumab versus 82.48% with nivolumab (**Figure 3C)**. While reducing TME’s protumoral bias and reprogramming M2 TAMs were both found to significantly reduce the M2:M1 ratio and increase CD8+ T cell counts, enhancing phagocytic activity of M1 TAMs has a direct effect on tumour cells, and thus causes a larger increase to the rate of treatment- and immune cell-induced tumour cell death than the first two strategies (**Figure 3D**). Indeed, the 0.01% maximal efficacy difference between the combination strategy with and without nivolumab suggests that solely targeting TAM phagocytic activity without an ICI may be sufficient to significantly increase glioblastoma survival.

### Treatment efficacy depends on M2:M1 TAM ratio

To understand whether enhancing phagocytic activity in M1 TAMs would remain efficacious a heterogeneous population, we ran a VCT on a cohort of virtual patients. Our global sensitivity analysis showed that the tumour burden after SOC and the supplementary efficacy of SOC+ICI were both statistically significantly sensitive to intrinsic tumour growth tumour and to the activity of the adaptive and innate immune systems (**Supplementary Figure 5**). Hence, we defined log-normal distributions for the intrinsic tumour growth rate, tumour cell antigenicity, and the TME bias towards the protumoral TAM phenotype (see **Virtual individual parameters** in **Supplementary Information**). We also sampled from previously estimated distributions^26^ for TMZ clearance, absorption, and plasma-to-CSF transfer rates to create 900 virtual patients (**Supplementary Figure 7**).

For each virtual patient (see Section **Generating a virtual cohort of patients and running a virtual clinical trial**), we modelled SOC with and without nivolumab. The addition of nivolumab was not predicted to benefit survival (**Supplementary Figure 8B**), consistent with previous clinical trial failures and our average model’s predictions. Unsurprisingly, patients with higher intrinsic tumour growth rates were those with the worst prognoses (**Supplementary Figure 8F**), consistent with our global sensitivity analysis (**Supplementary Figure 8A**). This underlines how the aggressive growth of glioblastomas results in high lethality, emphasizing the need to establish treatment strategies with a direct effect on tumour growth, such as enhancing M1 TAM phagocytic activity.

Thus, we modelled SOC, with or without nivolumab, in combination with an anti-CD47 antibody restoring glioblastoma cell phagocytosis by M1 TAMs in our virtual cohort. Our VCT predicted final tumour size decreases between 63.71% and 88.71% without PD-1 inhibition, with no additional benefit with nivolumab (**Figure 4A**). Survival was found to be significantly different (*p* < 10^−6^, log-rank test) between combination treatments that enhanced the phagocytic activity in M1 TAMs and those that did not (**Figure 4B** and **Supplementary Table 6**).

**Figure 4.**
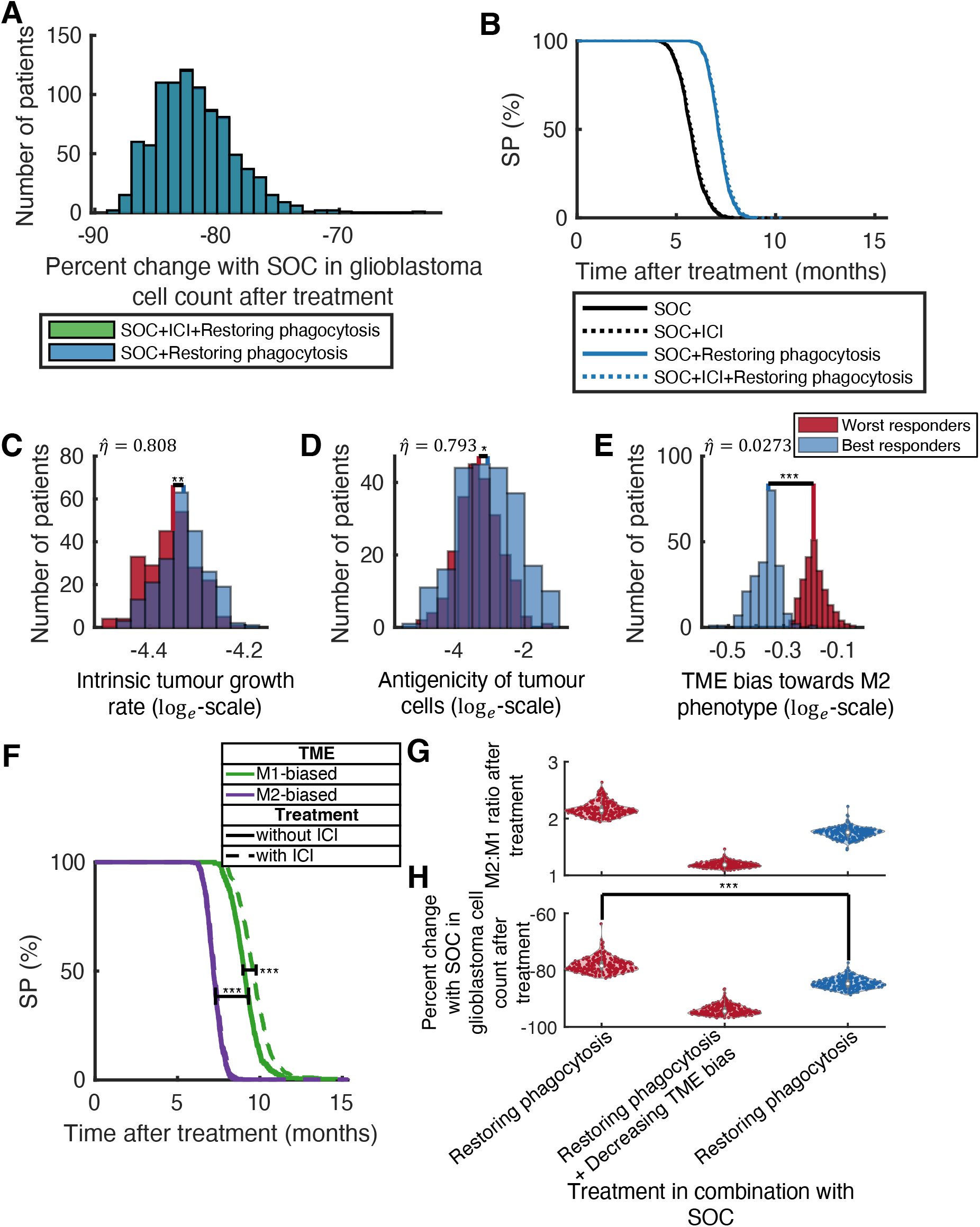
M2:M1 ratio is correlated with extra efficacy conferred by targeting TAMs. **A)** Percent change in glioblastoma cell count after treatment when M1 phagocytic activity is restored in combination with SOC (blue) and SOC+ICI (green). No difference is observed between the two scenarios, suggesting that restoring M1 phagocytic activity by targeting a “*don’t eat me signal*” such as CD47 in combination with SOC might be sufficient to improve prognoses. **B)** Kaplan-Meier survival curves for four combination treatment scenarios. Restoring M1 phagocytic activity significantly increases survival compared to other treatment scenarios. **C-D-E)** Histograms of the C) intrinsic tumour growth rate, D) antigenicity of tumour cells, and E) TME bias towards M2 phenotype for the worst (red) and best (blue) responders. In each, 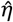 indicates the value of the overlapping index between distributions. TME bias towards M2 phenotype was the only patient-specific parameter found to distinguish best from worst responders. **F)** Kaplan-Meier (KM) curves for restoring M1 phagocytic activity in combination with SOC (solid) and SOC+ICI (dashed) in a M1- (green) and a M2- (purple) biased virtual populations. Survival was significantly different (*p* < 10^−6^, log-rank test) between M1- and M2-biased virtual populations for both combination scenarios. In the M1-biased virtual population, administering nivolumab significantly increased survival (*p* < 10^−6^, log-rank test), while it had no effect in the M2-biased virtual population. **G-H)** M2:M1 ratio and percent change with SOC in glioblastoma cell count after treatment (x-axis) in combination with SOC for the worst (red) and the best (blue) responders to the enhancement of phagocytosis of glioblastoma cells by M1 TAMs. ^*^: *p* < 10^−3, **^: *p* < 10^−6, ***^: *p* < 10^−16^ (numerical precision), two-sided Wilcoxon rank sum test for distributions and log-rank test for KM curves.

Next, we segregated the cohort into quartiles based on each virtual patient’s benefit, defined as the percent decrease in tumour cells after treatment compared to SOC normalized to the tumour intrinsic aggressiveness (see **Statistical analysis**. We defined the worst and best responders as the 25% of patients with the lowest and highest benefits, respectively. Our VCT predicted the percent decrease to be significantly higher in the best than in the worst responder group (**Figure 4H**, *p* < 10^−16^, two-sided Wilcoxon rank sum test). To identify which patient specific characteristics differentiated best from worst responders, we compared the distributions of the six parameters varied to generate the virtual population (see **Supplementary Table 7**). While best responders tended to have tumours with faster intrinsic growth and higher antigenicity than worst responders (*p* = 3.9014*e* − 06 and *p* = 0.0046 respectively), only the TME bias towards the M2 phenotype distinguished the best from worst responders (*p* < *e* − 16, 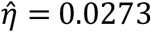). Additionally, the higher the TME bias towards the protumoral phenotype, the less a virtual patient was predicted to respond to the enhancement of M1 TAM phagocytic activity.

To further verify that the M2:M1 ratio predicts the benefit of combining a TAM-targeting treatment to SOC, we generated two new virtual cohorts: one with the fraction of TAMs acquiring the M1 phenotype upon activation log-normally distributed between 0 and 0.5 (M1-biased), and the other with the same fraction log-normally distributed between 0.5 and 1 (M2-biased; see **Supplementary Figures 9** and **10**). We simulated SOC in combination with anti-CD47, with or without nivolumab, in these virtual cohorts. Survival was found to be significantly higher in the M1-biased population (**Figure 4F**, *p* < 10^−16^, log-rank test) Notably, the addition of nivolumab to the SOC+anti-CD47 combination also significantly increased survival in this cohort (**Figure 4F**, *p* < 10^−16^, log-rank test).

We next asked whether reducing the M2:M1 ratio of the worst responders could increase the efficacy of phagocytosis enhancement in combination with SOC. Thus, in these virtual patients, we simulated SOC in combination with enhancing phagocytosis and increasing the M2-to-M1 repolarisation rate, since the latter strategy was shown to reduce the M2:M1 ratio (**Figure 3D**). In the worst responders, this combination increased treatment efficacy such that their percent decrease in tumour cell count after treatment was predicted to be larger than that of the best responders, who received no modulation of their M2:M1 ratio (**Figure 4H**). This suggests that patients with a TME highly biased towards M2-activated TAMs might benefit from combination TAM-targeting.

### Enhancing the phagocytic activity of antitumoral TAMs has significant potential in the clinic

Above, we assumed constant efficacy when simulating proposed TAM-targeting therapies. However, clinically relevant therapies will have time varying efficacy (i.e., pharmacodynamics (PDs)) determined by a drug’s pharmacokinetics (PKs). To study how the PK/PD of an anti-CD47 treatment to enhance the phagocytic activity of M1 TAMs affects predicted treatment outcomes, we modelled treatment efficacy to decay exponentially after administration, mimicking time-varying drug effects. For this, we considered a theoretical maximal effect of 1 and a half-life in the plasma of 13 days, based on the experimental molecule Hu5F9-G4, an anti-CD47 antibody investigated in a Phase 1 clinical trial for patients with advanced solid tumours^36^.

We then investigated weekly and biweekly administrations (**Figure 5A**). According to Swanson et al.^46^, a glioblastoma is considered to be detectable on enhanced CT scan at a volume equal to that of a sphere with a diameter of 3 cm, which is approximately equal to 14,137 · 10^6^ cells. We set this cell count as the detection threshold. For an average patient, we found that if anti-CD47 immunotherapy ended at the same time as SOC, glioblastoma cell counts one-week post-treatment in both administration scenarios were far below SOC (**Figure 5B**). More importantly, our model predicts that this final tumour volume would be under 2.5 cubic centimeters and thus undetectable, representing less than 15% of the size at diagnosis. However, given that treatment was ceased, all tumours were predicted to regrow to detectability eleven months after diagnosis. If the anti-CD47 immunotherapy was instead administered up to a year after the last TMZ administration in SOC, we found the tumour cell count to stabilize below a quarter of the lethal threshold, just above the detection threshold (**Figure 5B**). This suggests that enhancing tumour cell phagocytosis by antitumoral TAMs through CD47 may have significant clinical potential.

**Figure 5.**
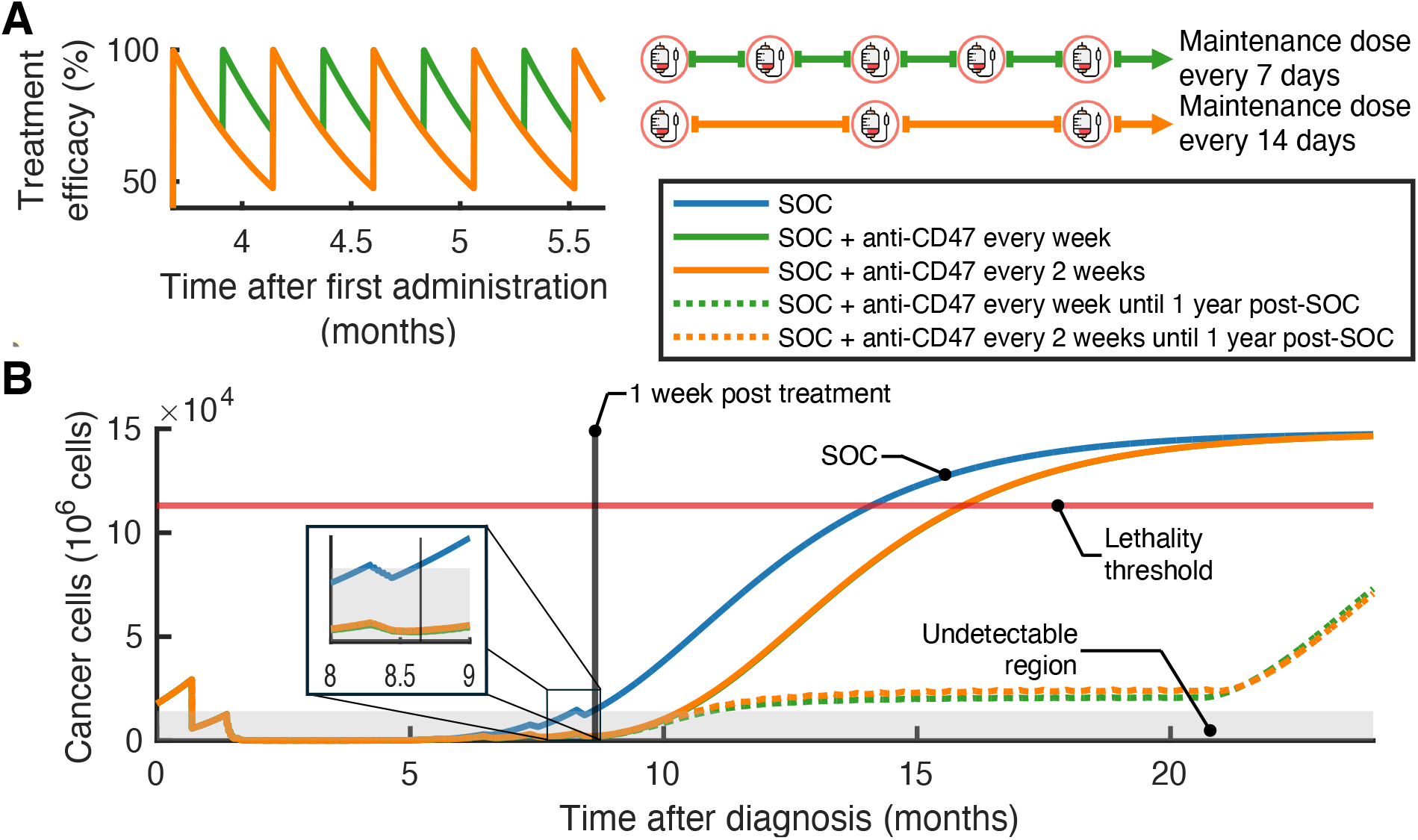
Anti-CD47 therapy in glioblastoma has significant clinical potential. **A)** To study how the PKs of a potential anti-CD47 molecule affect predicted treatment outcomes, TAM-targeting treatment efficacy was modelled to decay exponentially, with a half-life of 13 days and a maximal effect of 1. **B)** Model predictions of tumour size during different treatment administration scenarios. Blue: SOC alone. Green: SOC combined with a potential anti-CD47 agent administered every 7 days. Orange: SOC combined with a potential anti-CD47 agent administered every 14 days. TAM-targeting treatments were ceased at the same time as the last TMZ administration (solid lines) or one year post the last TMZ administration (dotted lines).

## Discussion

Despite advances in care, glioblastoma remains the most aggressive primary brain tumour, with a median survival of barely fifteen months^1,2^. To improve SOC, anti-PD-1 ICIs such as nivolumab have been intensely studied ^3^, with disappointing clinical trials results^5,7,29^. To better understand the mechanisms behind these failures, we developed a mathematical model describing both innate and adaptive immunity, and combination treatment with SOC and nivolumab that was validated to SOC+nivolumab glioblastoma clinical trial outcomes^5^. Our results show that nivolumab’s efficacy is limited by the lack of effector CD8+ T cells recruited to the TME during treatment. This explains why brain metastases, unlike glioblastomas, respond to nivolumab^47^, as they exhibit significantly higher CD8+ T cell counts than glioblastomas^18,41^. As such, our model’s predictions suggest that glioblastoma does not constitute a good candidate for anti-PD-1 ICIs unless administered in combination with a therapy that increases the recruitment of effector T cells to the TME.

Recent studies suggest that protumoral TAMs play an important role sustaining TME immunosuppression in glioblastomas^17^. The effective recruitment of CD8+ T cells is, in part, prevented by protumoral TAM polarisation. Thus, we investigated the potential of these tumour-supportive cells as clinical targets. We used our model to simulate five treatment strategies based on approaches currently undergoing clinical assessment. Our model predictions show that inhibiting TAM recruitment in glioblastoma TME via CCL2 inhibition, for example, does not significantly reduce post-treatment glioblastoma burden. Similarly, decreasing the survival of TAMs through CSF-1R blockade was also found to be ineffective. The use of anti-mouse CSF1R antagonist antibody (*α*CSF1R) in various tumour mouse models has been shown to significantly deplete *α*CSF1R-mediated anti- and protumoral TAMs. Hence, *α*CSF1R therapy fails to promote antitumor activity of the T cell compartment^31^. Our results suggest that this failure could be due to a lack of M2 target specificity.

Conversely, our model predicted that increasing TAM antitumoral functionalities could significantly reduce tumour burden. Maximally reactivating the phagocytotic activity of antitumoral TAMs by inhibiting “*don’t eat me*” signals was predicted to nearly completely eradicate the tumour. Using our model, we designed a VCT to recapitulate patient heterogeneity by generating 900 virtual patients with different TMZ pharmacokinetics profiles, intrinsic tumour growth rates, tumour cell antigenicity, and TME bias towards M2-activated TAMs. As we found no difference in median survival times with nivolumab in our VCT, our model predicts that targeting “*don’t eat me signals*” to enhance TAM phagocytotic activity could not only significantly increase the efficacy of SOC without an ICI. This could have potential ramifications for the toxicity of combination therapy. With our VCT framework, we found the M2:M1 ratio to be a predictor of the potential benefit of combining an anti-CD47 treatment with SOC. In particular, patients with a TME more strongly skewed towards M2 TAMs were predicted to benefit from the addition of a treatment repolarizing TAMs to the SOC+anti-CD47 combination. Based on these encouraging results, we simulated weekly and biweekly anti-CD47 mAB regimens, ending at or beyond the end of SOC, and showed that they have the potential to stabilize glioblastoma tumour size below a lethal size. These predictions agree with promising results from a first-in-human and first-in-class Phase I trial in patients with advanced two ovarian/fallopian tube cancers who achieved partial remission for over 5 months with the anti-CD47 antibody Hu5F9-G4^36^.

As our study was performed entirely *in silico*, follow-up experiments should be designed to validate our findings. For the sake of simplicity, we only included CD8+ T cells and TAMs as immune cells in our model and omitted other immune cell types. Neutrophils, for example, promote glioblastoma growth and enhanced neutrophil activity correlates with worse patient outcomes^48,49^. In hepatocellular carcinoma, tumour-associated neutrophils were shown to support tumour growth by promoting TAM infiltration^50^. However, this remains to be studied in glioblastoma. Our mathematical framework could be adapted in response to future experimental results quantifying neutrophil effects. Finally, although resistance was not included in our study, the Gompertz growth model we used implies that tumour growth rate tends to infinity as tumour cell count tends to 0^51^. Thus, our model always predicts rebound post-treatment, even if very few cells remain.

Taken together, this study shows that targeting TAMs by enhancing their antitumoral properties is a highly promising avenue to treat glioblastoma and reduce its lethality. As such, care should now be put on designing experiments to validate our findings. Here we provide proof-of-concept that mechanistic mathematical modelling and VCTs are complementary tools to uncover mechanisms driving clinical trial failures and explore the potential efficacy of novel treatment strategies by performing large-scale experimentation. Future work will also include adapting our model to promising drug candidates, to better understand their mechanism of action and optimise their administrations, helping to reduce attrition along the drug development pipeline and improve care for hard-to-treat tumours like glioblastoma.

## List of abbreviations

SOC: Standard-of-care
TMZ: Temozolomide
ICI: Immune checkpoint inhibitor
mAB: Monoclonal antibody
TME: Tumour microenvironment
TAM: Tumour-associated macrophages and microglia
TGF-β: Transforming growth factor beta
IL-10: Interleukin-10
TCGA: The Cancer Genome Atlas
VCT: Virtual clinical trial
SI: Supplementary information
CD47: Cluster of differentiation 47
CSF: Cerebrospinal fluid
CSF1R: Colony-stimulating factor-1 receptor
CCL2: Chemokine (C-C motif) ligand 2
TLR: Toll-like receptors

## Data and Code Availability Statement

The computational code to implement our model, and datasets resulting from simulations and needed for reproducing figures in the current study, are available on Github: https://github.com/Craig-Lab/TargetingTAMsGlioblastoma. Any further datasets are available from the corresponding authors upon reasonable request.

## Author’s Contributions

BM: Participated in research design, conducted experiments, contributed new analytic tools, performed data analysis, wrote the manuscript.

MC: Participated in research design, performed data analysis, wrote the manuscript, secured funding, supervised study.

## Acknowledgements

The authors wish to thank the institutions who funded this study. BM was funded by Fonds de recherche du Québec-Nature et Technologies, Natural Sciences and Research Council of Canada (NSERC) Master’s scholarships, an NSERC Canada Graduate Scholarship – Doctoral program (CGS D), and funding from the Université de Montréal. MC is the Canada Research Chair in Computational Immunology and this research was undertaken, in part, thanks to funding from the Canada Research Chairs Program. MC was also funded by an FRQS J1 Research Scholarship, the Fondation du CHU Sainte-Justine, and NSERC Discovery Grant RGPIN-2018-04546. Funders played no role in study design, data collection, analysis and interpretation of data, or the writing of this manuscript.

