## Supplementary information for "Virtual clinical trial reveals significant clinical potential of targeting tumour-associated macrophages and microglia to treat glioblastoma"

### Complete model equations

Our glioblastoma growth model describes the interactions between the tumour and its immune microenvironment by including CD8+T cells, resident and activated tumour-associated macrophages (TAMs), and PD-1 and PD-L1 receptors. We also track the amount of TMZ in the gastrointestinal tract, plasma, and cerebrospinal fluid to quantify TMZ concentration-dependant chemotherapy-induced glioblastoma cell killing. For the immune checkpoint inhibitor (ICI) model, we track the concentration in the tumour microenvironment of PD-1 receptors that are bound to an ICI. State variables of our model and their units are described in **Supplementary Table 1**.

| Variable | Description | Units |
|---|---|---|
| $G$ | Glioblastoma tumour cells | $10^6$ cells |
| $T$ | CD8+ T cells | $10^6$ cells |
| $M$ | Resident TAMs | $10^6$ cells |
| $M_1$ | M1-activated TAMs | $10^6$ cells |
| $M_2$ | M2-activated TAMs | $10^6$ cells |
| $PD1_U$ | Unbound PD-1 concentration in TME | pg/mL |
| $PD1_B$ | Bound PD-1 concentration in TME | pg/mL |
| $PDL1$ | PD-L1 concentration in TME | pg/mL |
| $Dd$ | Dead and damaged cells | $10^6$ cells |
| $I$ | ICI concentration in the TME | mg/mL |
| $A_1$ | Amount of TMZ in the gastrointestinal tract | mg |
| $A_2$ | Amount of TMZ in the plasma | mg |
| $A_3$ | Amount of TMZ in the cerebrospinal fluid (CSF) | mg |

**Supplementary Table 1. Variables of the mathematical model, with their description and units.**

The untreated glioblastoma growth and immune interactions model is given by:

$$\frac{dG}{dt} = \beta \cdot G \cdot \ln\left(\frac{K}{G}\right) - \alpha \frac{T}{T + k_1} G, \tag{S1}$$

$$\frac{dT}{dt} = -\delta_T \cdot T + \left( \alpha_{AT} \frac{G}{G + h_T} T + \eta_1 \frac{\frac{M_1}{M_1 + M_2}}{\eta_2 + \frac{M_2}{M_1 + M_2}} \right) \frac{1}{1 + \frac{PD1_U \cdot PDL1}{k_{YQ}}}, \tag{S2}$$

$$\frac{dM}{dt} = \lambda_M - a_{DM} \cdot M \cdot Dd - \delta_{MR} \cdot M - a_{MM} \cdot (M_1 + M_2) \cdot M, \tag{S3}$$

$$\frac{dM_1}{dt} = -\delta_M \cdot M_1 + (1-q) \cdot (a_{DM} \cdot M \cdot Dd + a_{MM} \cdot (M_1 + M_2) \cdot M) - \mu_{M1} \cdot M_1 + \mu_{M_2} \cdot M_2, \tag{S4}$$

$$\frac{dM_2}{dt} = -\delta_M \cdot M_2 + q \cdot (a_{DM} \cdot M \cdot Dd + a_{MM} \cdot (M_1 + M_2) \cdot M) - \mu_{M2} \cdot M_2 + \mu_{M_1} \cdot M_1, \tag{S5}$$

$$\frac{dPD1_U}{dt} = \rho_T \left( \alpha_{AT} \frac{G}{G + h_T} + \eta_1 \frac{\frac{M_1}{M_1 + M_2}}{\eta_2 + \frac{M_2}{M_1 + M_2}} \right) \frac{1}{1 + \frac{PD1_U \cdot PDL1}{k_{YQ}}} - \rho_T \cdot \delta_T \cdot T, \tag{S6}$$

$$\frac{dPDL1}{dt} = \rho_G \frac{dG}{dt} + \rho_M \left( \frac{dM}{dt} + \frac{dM_1}{dt} + \frac{dM_2}{dt} \right), \tag{S7}$$

$$\frac{dDd}{dt} = \alpha \frac{T}{T + k_1} G - \delta_{DM} \cdot Dd \cdot (M + M_1 + M_2) - d_{Dd} \cdot Dd. \tag{S8}$$

### Treatment with resection

SOC treatment of glioblastoma begins with surgical resection of the tumour. To model this, we used our glioblastoma growth model (Eqs. S1-S8) to predict tumour growth from diagnosis to the median time to surgical resection. Since we did not have data quantifying the resected proportions of all cell populations in our model, we simulated a direct reduction of all populations. Hence, let $r$ be the extent of the resection ($r < 1$). In a clinical trial conducted by Stupp et al. on patients with newly diagnosed, histologically confirmed glioblastomas[1], 126 and 113 patients receiving SOC went through partial and complete resections, respectively. Thus, we set $r = 0.8$ in our simulations to simulate a subtotal resection[2,3]. Then $C^+ = (1-r) \cdot C^-$, where $C^-$ is the value of population $C \in \{G, T, M, M_1, M_2, Dd, PD1_U, PDL1\}$ prior to resection and $C^+$ is its value post-resection.

### Treatment with radiotherapy

We used a direct tumour volume reduction[4] to model radiotherapy. Let $G_-$ and $G_+$ be the glioblastoma cell count prior to and after a radiation fraction, respectively, such that $G_+ = \gamma G_-$, where $\gamma$ is the cell survival probability. To define $\gamma$, we used the linear-quadratic (LQ) dose response model, which correlates the cell survival probability after a radiation fraction of size $D$ (Gy). In this model, $\gamma = \exp(-\alpha_{RT} D - \beta_{RT} D^2)$, where $\alpha_{RT}$ and $\beta_{RT}$ are two radiosensitivity parameters whose fraction $\alpha_{RT}/\beta_{RT}$ is tumoral-specific and describes the tumour's radiosensitivity[4–6]. Tumour cells damaged by radiotherapy become dead and damaged cells ($Dd$). During SOC, radiation is administered five days a week (2 Gy/day) concomitant to the six-week chemoradiotherapy cycle that follows surgical resection (see **Figure 2A** in the Main Text).

### Treatment with temozolomide

To describe TMZ concentrations in the body and at the tumour site over time, we integrated the three-compartment model population pharmacokinetic model developed by Ostermann et al.[7] (**Supplementary Figure 1A**). This model describes first-order absorption into the gastrointestinal tract (GIT) and linear elimination from the plasma:

$$\frac{dA_1}{dt} = -k_a A_1, \tag{S9}$$

$$\frac{dA_2}{dt} = k_a A_1 - \frac{CL}{V_D} A_2 - k_{23} A_2 + k_{32} A_3, \tag{S10}$$

$$\frac{dA_3}{dt} = k_{23}A_2 - k_{32}A_3, \tag{S11}$$

$$C_{plasma} = \frac{A_2}{V_D}, \tag{S12}$$

$$C_{CSF} = \frac{A_3}{V_p}. \tag{S13}$$

Here, $A_1$, $A_2$ and $A_3$ are the amount of TMZ in the GIT, plasma, and cerebrospinal fluid (CSF) compartments, respectively, while $C_{plasma}$ and $C_{CSF}$ are the drug concentrations in the plasma and the CSF compartments. $V_D$ and $V_p$ represent the TMZ volume of distribution in the central and CSF compartments, respectively, $k_a$ the absorption rate from the GIT, $k_{23}$ and $k_{32}$ the rates of exchange between the plasma and the CSF compartments, and $CL$ the plasma clearance.

We considered the CSF as the site of action, given that glioblastomas occur in the brain and central nervous system. We modelled TMZ pharmacodynamics using a standard Hill function:

$$E_{TMZ}=E_{max}\frac{C_{CSF}^h}{C_{CSF}^h+ IC50_{TMZ}^h}, \tag{S14}$$

where $E_{TMZ}$ denotes the anti-tumoral effect of the TMZ on glioblastoma cells, $E_{max}$ is the maximal effect, $h$ is the Hill coefficient describing the slope of the response curve, and $IC50_{TMZ}$ is the half maximal effective concentration, i.e., the TMZ concentration that produces an effect equal to $E_{max}/2$. Although TMZ is considered to be primarily cytostatic[8], we modelled its effects as cytotoxic, as our ODE model does not explicitly keep track of the growth rate of the tumour cells affected by TMZ. Hence, we assumed that temozolomide induces cell death in proliferating cells by arresting their cell cycle[9]. To model the effects of TMZ on glioblastoma cells, we replaced Eqs. (S1) and (S8) by:

$$\frac{dG}{dt}= \beta \cdot G \cdot \ln\left(\frac{K}{G}\right) - \alpha\frac{T}{T+k_1}G - E_{TMZ}G, \tag{S15}$$

$$\frac{dDd}{dt}=\alpha\frac{T}{T+k_1}G - \delta_{DM}\cdot Dd \cdot (M+M_1+M_2) - d_{Dd}\cdot Dd + E_{TMZ}G. \tag{S16}$$

### Immune checkpoint inhibitor dosing

Lee et al.[10] found that a nivolumab flat dosage of 240 mg every two weeks resulted in an exposure profile similar to that of a regimen of 3 mg/kg given once every two weeks. We assumed that nivolumab exhibits first order pharmacokinetics[11], implying that 480 mg and 6 mg/kg doses have similar exposures. Based on pharmacokinetic data from a phase 1 study by Brahmer et al.[12], Storey et al.[13] related the ICI dosage $D_{ICI}$ (mg/kg) to plasma concentration $C_{max}$ in $\mu$g/ml using

$$C_{max}(D_{ICI}) = 20D_{ICI} + 9.2. \tag{S17}$$

Then, converting the units of $C_{max}(D_{ICI})$ to mg/ml, we have that

$$C_{max}(D_{ICI}) = 0.02D + 0.0092. \tag{S18}$$

Nivolumab is administered as a one-hour intravenous infusion. Thus, we modelled the administration of the ICI as

$$I(t) = \begin{cases} 0.02D_{ICI} + 0.0092 & t_d \le t \le \frac{1}{24}. \\ 0 & \text{otherwise} \end{cases} \tag{S19}$$

where $t_d$ are the times of nivolumab administration and $D_{ICI}$ (mg/kg) is the dose.

Let $I(t)$ be the ICI plasma concentration in mg/ml. We modelled ICI dynamics according to:

$$\frac{dI}{dt} = I(t) - \mu_I PD1_U - \delta_I I, \tag{S20}$$

where the term $-\mu_I PD1_U$ represents the binding of nivolumab to unbound PD-1 receptors on CD8+ T cells and the rate $\delta_I$ (day$^{-1}$) represents its clearance from the body. We thus updated Eq. (S6) and integrated the following equations to specifically track unbound and bound anti-PD-1/PD-L1 concentrations:

$$\frac{dPD1_U}{dt} = \rho_T \left( \alpha_{AT} \frac{G}{G + h_T} + \eta_1 \frac{\frac{M_1}{M_1 + M_2}}{\eta_2 + \frac{M_2}{M_1 + M_2}} \right) \frac{1}{1 + \frac{PD1_U \cdot PDL1}{k_{YQ}}} - \frac{PD1_U}{PD1_U + PD1_B} \rho_T \cdot \delta_T \cdot T - \mu_P \cdot PD1_U \cdot I, \tag{S21}$$

$$\frac{dPD1_B}{dt} = -\frac{PD1_B}{PD1_U + PD1_B} \rho_T \cdot \delta_T \cdot T + \mu_P \cdot PD1_U \cdot I. \tag{S22}$$

Eq. (S21) describes the PD-1 receptors on the surface of CD8+ T cells that are bound to the anti-PD-1 and, thus, that cannot bind to PD-L1 on tumour cells and TAMs. The last terms in Eqs. (S21) and (S22) represent the binding of PD-1 receptors to the ICI. Since we do not know whether CD8+ T cells that die have bound or unbound receptors on their surface, we assumed that the amount of unbound (bound) PD-1 receptors removed from the TME is proportional to the percentage of PD-1 receptors that are unbound (bound). This is represented by the second term in Eq. (S21) and the first term in Eq. (S22).

### Targeting tumour-associated macrophages and microglia

To simulate TAM-targeting strategies, we introduced five parameters ($\lambda_{S1}, \lambda_{S2}, \lambda_{S3}, \lambda_{S4}$ and $\lambda_{S5}$) and modified Eqs. (S1), (S3), (S4), and (S5) to become:

$$\frac{dG}{dt} = \beta \cdot G \cdot \ln\left(\frac{K}{G}\right) - \alpha \frac{T}{T + k_1} G - E_{TMZ} G - \lambda_{S3} \cdot d_{M_{\phi_1} G} \cdot G \cdot M_1, \tag{S23}$$

$$\frac{dM}{dt} = \lambda_{S2} \cdot \lambda_M - a_{DM} \cdot M \cdot Dd - \lambda_{S1} \cdot \delta_{MR} \cdot M - a_{MM} \cdot (M_1 + M_2) \cdot M, \tag{S24}$$

$$\frac{dM_1}{dt} = -\lambda_{S1} \cdot \delta_M \cdot M_1 + (1 - \lambda_{S5} \cdot q) \cdot (a_{DM} \cdot M \cdot Dd + a_{MM} \cdot (M_1 + M_2) \cdot M) - \mu_{M1} \cdot M_1 + \lambda_{S4} \cdot \mu_{M_2} \cdot M_2, \tag{S25}$$

$$\frac{dM_2}{dt} = -\lambda_{S1} \cdot \delta_M \cdot M_2 + \lambda_{S5} \cdot q \cdot (a_{DM} \cdot M \cdot Dd + a_{MM} \cdot (M_1 + M_2) \cdot M) - \lambda_{S4} \cdot \mu_{M2} \cdot M_2 + \mu_{M_1} \cdot M_1, \tag{S26}$$

Each lambda-term is defined by

$$\lambda = \begin{cases} \lambda_0, & \text{off treatment} \\ s, & \text{during treatment} \end{cases}.$$

Off treatment values and intervals of definition for $s$ during treatment are given in **Supplementary Table 2**. The value of $s$ during treatment determines its efficacy: the closer $s$ is to $\lambda_0$, the less effective the simulated treatment, while the other end of its interval of definition represents maximal efficacy.

| TAM-targeting strategy | TAM-targeting parameter | Off treatment value ($\lambda_0$) | Interval of definition for $s$ |
|---|---|---|---|
| Delete TAMs | $\lambda_{S1}$ | 1 | $\left(1, \frac{1}{\max(\delta_M, \delta_{MR})}\right)$ |
| Inhibit of TAMs | $\lambda_{S2}$ | 1 | (0,1) |
| Restore phagocytosis of cancer cells by TAMs | $\lambda_{S3}$ | 0 | (0,1] |
| Reprogram TAMs into TAMs with anti-tumoural properties | $\lambda_{S4}$ | 1 | $\left(1, \frac{1}{\mu_{M2}}\right)$ |
| | $\lambda_{S5}$ | 1 | (0,1) |

**Supplementary Table 2. Parameter definitions for simulating TAM-targeting strategies**

$\lambda_{S1}$ and $\lambda_{S2}$ were used to model TAM-targeting strategies that reduce the total number of TAMs in the TME. $\lambda_{S1}$ simulates TAM depletion by increasing the natural death rate of TAMs, whereas $\lambda_{S2}$ models the inhibition of TAM recruitment to the TME by decreasing the constant source of TAMs ($\lambda_M$). We assumed that the macrophage pool could not turnover more than once per day (i.e., $\lambda_{S1} \cdot \delta_M$ for M1 and M2 macrophages and $\lambda_{S1} \cdot \delta_{MR}$ for tissue-resident TAMs). This bounds $\lambda_{S1}$ above by $\frac{1}{\max(\delta_M, \delta_{MR})}$.

$\lambda_{S3}$ restores the phagocytosis of tumour cells by anti-tumoral TAMs. We assumed that without treatment, tumour cells could completely escape TAM phagocytosis. $\lambda_{S4}$ and $\lambda_{S5}$ both simulate TME reprogramming into an anti-tumoral environment through two different routes. $\lambda_{S4}$ confers an M1-like phenotype to already activated pro-tumoral TAMs by increasing the rate M2 TAMs switch to a M1 phenotype. Whereas $\lambda_{S5}$ decreases the fraction of resident TAMs that acquire a pro-tumoral phenotype upon activation. During treatment, $\lambda_{S4}$ is bounded by above by $\frac{1}{\mu_{M2}}$, as we assumed, as above, that the total daily population turnover rate for macrophages to be 100%. The range spanned by $\lambda_{S5}$ models treatment efficacies between no decrease in the TME bias towards the M2 phenotype ($\lambda_{S5} = 1$) and no newly activated TAMs acquiring the M2 phenotype ($\lambda_{S5} = 0$).

### Predicting survival time

Based on a cohort of 106 patients diagnosed with histopathologically-verified glioblastomas, Stensjøen et al.[14] determined the median tumour volume at diagnosis to be 17.7 mL. As water accounts for more than 70% of the total mass of a cell[15], we took 1 mL=1 $cm^3$, giving a median tumour volume of 17.7 $cm^3$. Assuming that human tumours contain about $10^9$ cells per cubic cm[16], this implies that the median cell count at diagnosis of $C_{diagnosis} = 17.7 \cdot 10^9$ cells. Swanson et al.[17] analysed observations of living and dead patients diagnosed with glioblastoma and found that the fatal average tumour volume to be equal to that of a sphere with a radius of 3 cm. Together, this gives us a fatal cell count threshold of about $C_{death} = 113.1 \cdot 10^9$ cells. Thus, we modelled survival time as the time it took to the cancer cell count to go from $C_{diagnosis}$ to $C_{death}$[17].

### Detailed parameter estimation calculations

We estimated the parameters in our model by 1) directly sourcing them from the literature, 2) fitting them to publicly available data, or 3) calculating them at homeostasis in the absence of a tumour. The glioblastoma cell carrying capacity was estimated by Stensjøen et al.[14] using data from 106 patients with glioblastoma. We then rescaled their value to our units. For the tumour-immune system model, the activation rate of CD8+ T cells by glioblastoma cells ($\alpha_{AT}$) was taken from Mahasa et al.[18]. From their model, we also calculated the half-saturation constant of the cytotoxic activity of the adaptive immune system ($k_1$). The inhibition constant of the CD8+ T cells activation by the PD-1/PD-L1 signalling axis $\left(k_{YQ}\right)$ was taken from Lai and Friedman[19] and rescaled to our units. The death rates of resident TAMs ($\delta_{MR}$) and activated TAMs ($\delta_M$) were calculated from the cells' half-lives. The death rate of primed CD8+ T cells was estimated by Kim et al.[20], while the phagocytosis parameters ($a_{DM}$ and $\delta_{DM}$) were taken and rescaled from Jenner et al.[21] As in Reynolds et al.[22], we assumed that resident TAMs get activated by $M_1$ and $M_2$ TAMs at a rate $a_{MM}$ that is ten times less the rate they get activated by phagocytes.

The natural clearance rate of dead and damaged cells ($d_{Dd}$) was taken from Jenner et al.[21] who directly estimated it from Elmore[23]. The switching rates from one TAM phenotype to another ($\mu_{M1}$ and $\mu_{M2}$) were taken from Wang et al.[24] The half-saturation constant of CD8+ T cell activation by the innate immune system ($\eta_2$) was taken from Louzoun et al.[25]. The PD-1 concentration per $10^6$ CD8+ T cells ($\rho_T$), and the PD-L1 concentration per $10^6$ tumour cells and TAMs ($\rho_G$ and $\rho_M$) are calculated from the number of PD-1 and PD-L1 receptors CD8+ T cells, assuming that there is ten times more PD-L1 receptors on tumour cells and TAMs than on CD8+ T cells[13]. Full details of the calculations are provided below and parameter values are given in **Supplementary Tables Supplementary** Table 2-**Supplementary** Table 10.

**1. <u>Parameters estimated from literature</u>**

1.1. <u>Tumour microenvironment (TME) volume</u>

Considering a carrying capacity (in density) of $0.8\,\frac{g}{cm^3}$, we estimated the total TME volume (in $cm^3$) as in Storey et al.[13] per the equation:

$$V = \frac{158040 \cdot 10^6 cells}{0.8\,\frac{g}{cm^3} \cdot 10^9\,\frac{cells}{g}} = 197.55 cm^3 = 197.55 ml$$

1.2. <u>Tumour carrying capacity ($K$)</u>

Stensjøen et al.[26] fit a Gompertz growth curve to data from 106 patients with glioblastoma and found a carrying capacity of $K$=158.04 ml. Again assuming that 1 ml = 1 $cm^3$ and that human tumours contain about $10^9$ cells per cubic cm[16], we obtained

$$K = 158.04\text{cm}^3 \cdot \frac{10^9\text{cells}}{\text{cm}^3} \cdot \frac{1000 \cdot 10^6\text{cells}}{10^9\text{cells}} = 158\,040 \cdot 10^6\text{cells}.$$

1.3. <u>Half-saturation constant of CD8+ T cell cytotoxic activity ($k_1$)</u>

Mahasa et al.[18] used $k_1 = 40$ cells for an initial CD8+ T cell population size of zero. We rescaled this value by multiplying it by the initial number of CD8+ T cells in our model (previously estimated from glioblastoma biopsy and resection imaging mass cytometry data[27,28]):

$$k_1 = 40 \cdot T(0) = 40 \cdot 0.0320 \cdot 10^6 cells = 1.28 \cdot 10^6\ cells$$

1.4. <u>Phagocytosis rate of glioblastoma cells by M1 TAMs $\left(d_{M_{\Phi 1}G}\right)$</u>

From Baba et al.[29], Shu et al.[30] estimated the killing rate of tumour cells by M1 cells to be $f = 2 \times 10^{-6} cells^{-1} day^{-1}$. Given $T$ and $M_1$ the tumour cells (cells) and M1 TAMs (cells), respectively, the rate of tumour cell clearance by M1 macrophages is of the form $-fTM_1$. In our model, tumour cells and TAMs are in units of $10^6$ cells. Thus, $d_{M_{\Phi 1}G}$ should be expressed in units of $\frac{1}{10^6 cells \cdot day}$, hence

$$d_{M_{\phi 1}G} = 2 \cdot 10^{-6} \frac{1}{cells \cdot day} \times 10^6 \frac{cells}{10^6 cells} = 2 \frac{1}{10^6 cells \cdot day}.$$

1.5. Half-saturation constant of CD8+ T cell activation by glioblastoma cells ($h_T$)

Mahasa et al.[18] reported the half-saturation constant for tumour cells in response to tumour antigens to be 40 cells. We calibrated this to our carrying capacity as:

$$\frac{40\ cells}{10^{11} cells} = \frac{h_T}{158040 \cdot 10^6 cells} \Rightarrow h_T = 6.3216 \cdot 10^{-5} \cdot 10^6 cells.$$

1.6. Inhibition of the CD8+ T cell activation by the PD-1/PD-L1 signalling axis ($k_{YQ}$)

Lai and Friedman[19] estimated $k_{YQ}$ to be $1.365 \cdot 10^{-18} \frac{g^2}{cm^6}$. We converted this to our units as

$$k_{YQ} = 1.365 \cdot 10^{-18} \frac{g^2}{cm^6} \cdot 10^{24} \frac{pg^2}{g^2} \cdot 1 \frac{cm^6}{ml^2} = 1.365 \cdot 10^6 \frac{pg^2}{ml^2}.$$

1.7. Activation rate of TAMs by dead and damaged cancer cells ($a_{DM}$)

Jenner et al.[21] estimated the rate of macrophage activation following an interaction with dead cells to be $1.1 \cdot 10^3 \frac{ml}{10^9 cells \cdot day}$, which when converted to the units in our model gives

$$a_{DM} = 1.1 \cdot 10^3 \frac{ml}{10^9 cells \cdot day} \cdot 10^{-3} \frac{10^9 cells}{10^6 cells} \cdot \frac{1}{197.55} \frac{1}{ml}$$
$$= 0.005568 \frac{1}{10^6 cells \cdot day}.$$

We assumed all TAMs to have the same activation rate as the macrophages modelled by Jenner et al. [21].

1.8. Death rate of resident TAMs ($\delta_{MR}$)

Mature macrophages have a half-life of 4-6 weeks[31], which translates to a death rate of

$$\delta_{MR} = \frac{ln(2)}{5 \text{ weeks} \cdot \frac{7 \text{ days}}{\text{week}}} = \frac{0.0198}{\text{day}}.$$

We assumed that resident TAMs have the same death rate as mature macrophages.

1.9. Death rate of M1 and M2 TAMs ($\delta_M$)

Macrophages have a half-life of approximately 3.4 days[32,33] and hence a death rate of

$$\delta_{MR} = \frac{ln(2)}{3.4 \text{ days}} = \frac{0.02}{\text{day}}.$$

We assumed all M1 and M2 TAMs to have the same death rate as macrophages.

1.10. Activation rate of resident TAMs by M1and M2 TAMs ($a_{MM}$)

Reynolds et al.[22] reported that previously activated immune cells are reactivated at a rate that is 0.1 time the rate of activation by phagocytes. Since in our model $a_{DM} = 0.0073 \frac{1}{10^6 \text{ cells} \cdot \text{ day}}$, this gives

$$a_{MM} = 0.1 \cdot 0.005568 \frac{1}{10^6 \text{ cells} \cdot \text{ day}} = \frac{0.0005568}{10^6 \text{ cells} \cdot \text{ day}}.$$

1.11. TME bias towards M2-activated TAMs ($q$)

We chose $q$ according to the reported M2:M1 ratio of 3.2 in glioblastoma patients[34]. Since $q$ is the fraction of resident TAMs that are activated to a M2 state, we have that

$$\frac{M_2}{M_1 + M_2} = \frac{3.2 \cdot M_1}{M_1 + 3.2 \cdot M_1} = \frac{16}{21}.$$

1.12. PD-L1 concentration per $10^6$ T cells ($\rho_T$)

Storey et al.[13] reported that there are 3096 PD-1 proteins expressed by each T cell, and that one PD-L1 protein weighs $8.3 \cdot 10^{-20}$g[35,36]. Hence, the PD-1 concentration per $10^6$ T cells (pg/ml) was calculated as:

$$\rho_T = 3096 \frac{PD - 1\, protein}{T\, cell} \cdot 8.3 \cdot 10^{-20} \frac{g}{PD - L1\, protein} \cdot \frac{1}{197.55} \frac{1}{ml} \cdot 10^6 \frac{T\, cells}{10^6\, T\, cells} \cdot 10^{12} \frac{pg}{g} = 1.3008 \frac{pg}{ml \cdot 10^6\, T\, cells}.$$

1.13. PD-L1 concentration per $10^6$ tumour cells ($\rho_G$) and $10^6$ TAMs ($\rho_M$)

As in Storey et al.[13], we assumed $\rho_G = \rho_M$. Given that there are 9282 PD-L1 proteins expressed on each T cell[13] and that one PD-L1 protein weighs $5.8 \cdot 10^{-20}$ g[36], the PD-L1 concentration per $10^6$ T cells in pg/ml was calculated to be

$$9282 \frac{PD - L1\, protein}{T\, cell} \cdot 5.8 \cdot 10^{-20} \frac{g}{PD - L1\, protein} \cdot \frac{1}{197.55} \frac{1}{ml} \cdot 10^6 \frac{T\, cells}{10^6\, T\, cells} \cdot 10^{12} \frac{pg}{g} = 2.72516 \frac{pg}{ml \cdot 10^6\, T\, cells}.$$

Storey et al.[13] also reported that the PD-L1 concentration per tumour cell and per TAM is 10 times the PD-L1 concentration per T cell. Hence, the concentration of PD-L1 per $10^6$ tumour cells and $10^6$ TAMs were calculated to be

$$\rho_G = \rho_M = 10 \cdot 2.72516 \frac{pg}{ml \cdot 10^6 Tcells} = 27.2516 \frac{pg}{ml \cdot 10^6 cells}.$$

1.14. Phagocytosis rate of dead and damaged glioblastoma cells by TAMs ($\delta_{DM}$)

We estimated $\delta_{DM}$ directly from Jenner et al.[21]

$$\delta_{DM} = 8.03 \frac{ml}{10^9 cells \cdot day} \cdot 10^{-3} \frac{10^9 cells}{10^6 cells} \cdot \frac{1}{197.55} \frac{1}{ml} = 4.0648 \cdot 10^{-5} \frac{1}{10^6 cells \cdot day}.$$

1.15. Binding rate of anti-PD-1 to PD-1 receptors ($\mu_I$)

We based our calculations of binding rates upon those in Storey et al.[13]. As in Lai and Friedman[19] and Storey et al.[13], we assumed that 10% of the ICI is used to block PD-1 receptors while the remaining 90% degrades naturally, giving

$$\frac{\mu_I \cdot P \cdot I}{0.1} = \frac{\delta_I \cdot I}{0.9}.$$

At steady state $\bar{P}$, we assumed

$$\mu_I = \frac{\delta_I}{9 \cdot \bar{P}}.$$

As the PD-1 carrying capacity is of $7.47 \cdot 10^{-7} \frac{\text{g}}{\text{L}}$, we found

$$\bar{P} = 7.47 \cdot 10^{-7} \frac{\text{g}}{\text{L}} \cdot 10^{12} \frac{\text{pg}}{\text{g}} \cdot \frac{1}{10^3} \frac{\text{L}}{\text{ml}} = 7.47 \cdot 10^2 \frac{\text{pg}}{\text{ml}},$$

$$\mu_I = \frac{\frac{0.0257}{\text{day}}}{9 \cdot 7.47 \cdot 10^2 \frac{\text{pg}}{\text{ml}}} = 3.8186 \cdot 10^{-6} \frac{\text{ml}}{\text{pg} \cdot \text{day}}.$$

1.16. Anti-PD-1 blocking rate of PD-1 receptors ($\mu_P$)

In our model, $\mu_P$ is expressed in units $\frac{\text{ml}}{\text{mg}\cdot\text{day}}$. We used the conversion

$$\mu_P = \mu_I \cdot 10^9 \frac{\text{pg}}{\text{mg}} = 3.8186 \cdot 10^3 \frac{\text{ml}}{\text{mg} \cdot \text{day}}.$$

1.17. Nivolumab clearance rate ($\delta_I$)

Nivolumab has a half-life of 27 days[13,37], implying

$$\delta_I = \frac{ln(2)}{27} = \frac{0.0257}{\text{day}}.$$

### 2. Parameters fitted to or calculated from publicly available data

2.1. Radiotherapy linear-quadratic parameters ($\alpha_{RT}$ and $\beta_{RT}$)

Franken et al.[38] estimated the linear quadratic parameters of glioma cell lines AMC-3046, VU-122 and VU-109 after exposure to TMZ-radiation treatment. Since glioblastoma tumours behave like early responding tissue when given a radiation fraction[39], this gives[40]

$$\frac{\alpha_{RT}}{\beta_{RT}} = 10 \implies \beta_{RT} = \frac{\alpha_{RT}}{10}.$$

We used parameter values for the VU-109 cell line as it was the one with the $\frac{\alpha}{\beta}$ ratio closest to 10 of the three cell lines. This gives $\alpha_{RT} = 0.19$ and $\beta_{RT} = 0.032$ $\left(\frac{\alpha_{RT}}{\beta_{RT}} = 6\right)$.

2.2. TMZ pharmacodynamic model

TMZ pharmacodynamics were described using the standard saturating dose-resposne Hill function[27,41–43] per Eq. (S14). Lo Dico et al.[44] measured dose-response curves for responsive glioblastoma cells after temozolomide treatment. We used U87 glioblastoma cell line viability after 72 hours of treatment at increasing doses of TMZ under normoxic conditions. We converted the Lo Dico et al. concentration data from $\mu M$ to mg/ml using

$$C_2 = C_1 \cdot \frac{10^{-6}\text{mol}}{\mu\text{mol}} \cdot \frac{L}{1000\text{mL}} \cdot 194.151 \frac{\text{g}}{\text{mol}} \cdot \frac{1000\text{ mg}}{\text{g}},$$

with $C_1$ in µM and $C_2$ in mg/mL, and 194.151I g/mol the molar weight of TMZ[45]. We then fit Eq. ((S14) using the function *lsqnonlin* in MatlabR2023b[46] which solves nonlinear least-square curve fitting problems and returns a set of parameter values minimizing

$$\sum_{i=1}^{n} \left(\hat{y}_i - y_i\right)^2, \tag{S27}$$

where $\hat{y}_i$ is the $i$-th data point from Lo Dico et al.'s cell viability assays[44] and $y_i$ is the cytotoxic effect with an equivalent TMZ concentration predicted with Eq. (S14). Since the data were reported as cell viability after 72 hours, we assumed that $E = 1 -$ cell viability and that $E_{max} = E_{max_{72\text{ hours}}}/3$, where $E_{max_{72\text{ hours}}}$ is the maximal effect obtained through fitting. This gave $E_{max} = 0.30010$, $IC50_{TMZ} = 0.0011 \frac{\text{mg}}{\text{ml}}$, and $h = 0.5682$. Results of the fits are shown on **Supplementary Figure 1**.

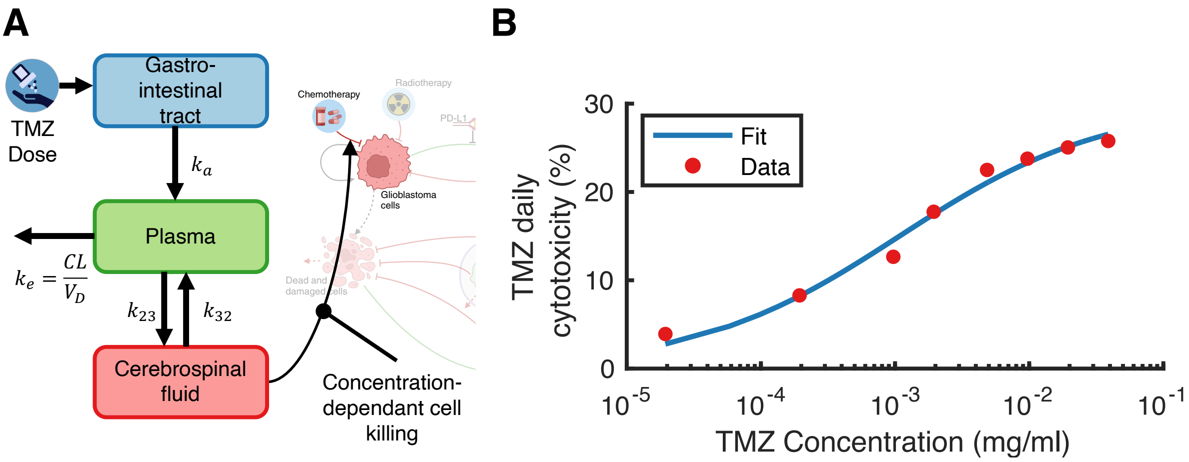


**Supplementary Figure 1. TMZ pharmacokinetics and pharmacodynamics. A)** PK/PD model schematic. We integrated the three-compartment population pharmacokinetic model described in Ostermann et al.[7] with first-order absorption in the gastrointestinal tract (GIT) and linear plasma elimination. Equations are detailed in the section **Treatment with temozolomide** in the Main Text. TMZ was modeled as a cytotoxic drug with a cerebrospinal fluid (CSF) concentration-dependant cell killing. **B)** The parameters of the TMZ dose-response relationship were parameterized by fitting the Hill equation in (Eq. (S14)) to data from cell viability assays performed by Lo Dico et al. on U87 GBM cells[44].

2.3. Intrinsic tumour growth rate ($\beta$) and glioblastoma killing rate by CD8+ T cells ($\alpha$)

The median survival time of an untreated glioblastoma is about 4 months[47]. Stupp et al.[1] reported a median survival time of 12.1 months after treatment with radiotherapy and of 14.6 months with combination radiotherapy and temozolomide. Based on these values, we calibrated $\beta$ and $\alpha$ with the function *lsqnonlin* in MatlabR2023b[46] to minimize Eq. (S27), where $\hat{y}_i$ is the observed survival time with no treatment ($i = 1$), RT ($i = 2$), and RT+TMZ ($i = 3$), while $y_i$ is the corresponding survival time predicted with the model. This fitting resulted in $\beta = 0.013$ and $\alpha = 0.027$, which means that T cells kill at most 2.7% of the tumour per day. Results of both Surgery+RT and SOC were found to fall within 95% confidence intervals. However, our model predicted a survival time without treatment of slightly over 5 months (**Supplementary Figure 2**). In our model, we used a constant volume at diagnosis. However, if a patient receives no treatment, the tumour of a patient not receiving treatment may have been diagnosed later, with a larger volume at diagnosis thus reducing our simulated survival time. Hence, we accepted the discrepancy of about 1 month.

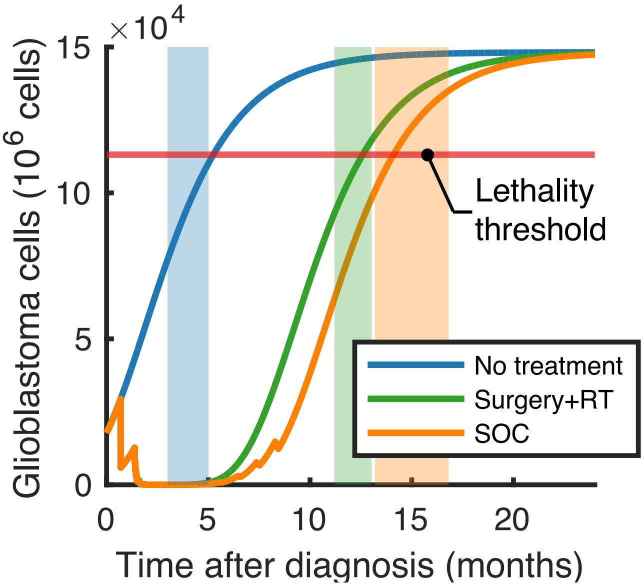


**Supplementary Figure 2. Estimating intrinsic tumour growth rate and CD8+ T cell killing rate from survival data.** The median survival time of an untreated glioblastoma is 4 months. Blue shaded region: median survival without treatment plus and minus one month. Green shaded region: median survival with resecetion + RT. Survival was predicted to increase to 12.1 months (95% C.I.: [11.2 13]). Orange shaded region: median survival after SOC. Our model predicted survival of 14.1 months (95% C.I.: [13.2 16.8]). Solid lines: model predictions.

### 3. Parameters calculated at no tumour homeostasis

We set homeostatic concentrations in absence of a tumour as the initial conditions in our model.

#### 3.1. Maximal activation rate of CD8+ T cells by TAMs ($\eta_1$)

Setting $\frac{dT}{dt} = 0$ with $M^* = M(0)$, $M_1^* = M_1(0)$, $M_2^* = M_2(0)$, $T^* = T(0)$, $P^* = \rho_P \cdot T^*$ and $D^* = \rho_M \cdot (M_1^* + M_2^* + M^*)$ in the absence of a tumour, we have:

$$0 = -\delta_T \cdot T^* + \left( \eta_1 \frac{\frac{M_1^*}{M_1^* + M_2^*}}{\eta_2 + \frac{M_2^*}{M_1^* + M_2^*}} \right) \cdot \left( \frac{1}{1 + \frac{P^* \cdot D^*}{k_{YQ}}} \right) \Rightarrow \eta_1 = 0.046336.$$

#### 3.2. Constant source of resident TAMs in the TME ($\lambda_M$)

At the no tumour equilibrium,

$$\frac{dM}{dt} + \frac{dM_1}{dt} + \frac{dM_2}{dt} = 0 \implies \lambda_M = \delta_M (M_1^* + M_2^*) + \delta_{MR} \cdot M^* = 0.0248 \frac{10^6 \text{cells}}{\text{day}}.$$

| Parameter | Definition | Units | Estimated value | Reference |
|---|---|---|---|---|
| $\beta$ | Tumour intrinsic growth rate | $\frac{1}{\text{day}}$ | 0.013 | Fit |
| $K$ | Carrying capacity of the cancer cells | $10^6$ cells | 158 040 | Calculated[26] |
| $\alpha$ | Maximal killing rate of the GBM cells by the CD8+ T cells | $\frac{1}{\text{day}}$ | 0.027 | Fit |
| $k_1$ | Half-saturating constant of the | $10^6$ cells | 1.28 | Calculated[18] |

| | | | | |
|---|---|---|---|---|
| | cytotoxic activity of the CD8+ T cells | | | |
| $d_{M_{\phi_1}G}$ | Phagocytosis rate of GBM by M1 TAMs | $\frac{1}{\text{day} \cdot 10^6\text{cells}}$ | 2 | Calculated[30] |

**Supplementary Table 3. Parameters of the cancer cell equation.**

| Parameter | Definition | Units | Estimated value | Reference |
|---|---|---|---|---|
| $\delta_T$ | Natural death rate of the CD8+ T cells | $\frac{1}{\text{day}}$ | 0.4 | Fixed[20] |
| $\alpha_{AT}$ | Maximal activation rate of the CD8+ T cells by the GBM cells | $\frac{1}{\text{day}}$ | 0.0375 | Fixed[18] |
| $h_T$ | Half-saturating constant of the activation of the CD8+ T cells by the GBM cells | $10^6\text{cells}$ | $6.3216 \cdot 10^{-5}$ | Calculated[18] |
| $\eta_1$ | Maximal activation rate of the CD8+ T cells by the innate immune system | $\frac{10^6\text{cells}}{\text{day}}$ | 0.0464 | Calculated at no tumour homeostasis |
| $\eta_2$ | Half-saturating constant of the activation of the CD8+ T cells by the innate immune system | Unit-less | 0.1 | Fixed[25] |
| $k_{YQ}$ | Inhibition of the CD8+ T cell activation by the PD-1/PD-L1 signalling axis | $\frac{\text{pg}^2}{\text{ml}^2}$ | $1.365 \cdot 10^6$ | Calculated[19] |

**Supplementary Table 4. Parameters of the CD8+ T cell equation.**

| Parameter | Definition | Units | Estimated value | Reference |
|---|---|---|---|---|
| $\lambda_M$ | Recruitment rate of resident TAMs to the TME | $\frac{10^6\text{cells}}{\text{day}}$ | 0.0248 | Calculated at no tumour homeostasis |
| $a_{DM}$ | Activation rate of resident TAMs by dead and damaged GBM cells | $\frac{1}{10^6\text{cells} \cdot \text{day}}$ | 0.0056 | Calculated[21] |
| $\delta_{MR}$ | Natural death rate of resident TAMs | $\frac{1}{\text{day}}$ | 0.0198 | Calculated[31] |
| $a_{MM}$ | Activation rate of resident TAMs by M1 and M2 activated TAMs | $\frac{1}{10^6\text{cells} \cdot \text{day}}$ | $5.5683 \cdot 10^{-4}$ | Calculated[21,22] |
| $\delta_M$ | Natural death rate of activated M1 and M2 TAMs | $\frac{1}{\text{day}}$ | 0.2 | Calculated[32,33] |
| $q$ | Fraction of resident TAMs that first get the anti-tumoral phenotype once activated | Unit-less | $\frac{16}{21}$ | Calculated[34] |

| | | | | |
|---|---|---|---|---|
| $\mu_{M1}$ | M1 → M2 repolarization rate | $\frac{1}{day}$ | 0.075 | Fixed[24] |
| $\mu_{M2}$ | M2 → M1 repolarization rate | $\frac{1}{day}$ | 0.05 | Fixed[24,48] |

**Supplementary Table 5. Parameters of the TAM equations ($M$, $M_1$ and $M_2$).**

| Parameter | Definition | Units | Estimated value | Reference |
|---|---|---|---|---|
| $\rho_T$ | PD-1 concentration per $10^6$ CD8+ T cells | $\frac{\frac{pg}{ml}}{10^6 cells}$ | 1.3008 | Calculated[13,35,36] |
| $\rho_G$ | PD-1 concentration per $10^6$ cancer cells | $\frac{\frac{pg}{ml}}{10^6 cells}$ | 27.2516 | Calculated[13,36] |
| $\rho_M$ | PD-1 concentration per $10^6$ TAMs cells | $\frac{\frac{pg}{ml}}{10^6 cells}$ | 27.2516 | Calculated[13,36] |

**Supplementary Table 6. Parameters of the PD-1 and PD-L1 equations.**

| Parameter | Definition | Units | Estimated value | Reference |
|---|---|---|---|---|
| $\delta_{DM}$ | Phagocytosis rate of dead and damaged cells by TAMs | $\frac{1}{10^6 cells \cdot day}$ | $4.0648 \cdot 10^{-5}$ | Calculated[21] |
| $d_{Dd}$ | Natural clearance of dead and damaged cells | $\frac{1}{day}$ | 8 | Fixed[21,23] |

**Supplementary Table 7. Parameters of the dead and damaged cell equation.**

| Parameter | Definition | Units | Estimated value | Reference |
|---|---|---|---|---|
| $\alpha_{RT}$ | Radiosensitivity parameter | $\frac{1}{Gy}$ | 0.019 | Fixed[38–40] |
| $\beta_{RT}$ | Radiosensitivity parameter | $\frac{1}{Gy^2}$ | 0.032 | Fixed[38–40] |

**Supplementary Table 8. Parameters of the radiotherapy treatment.**

| Parameter | Definition | Units | Estimated value | Reference |
|---|---|---|---|---|
| $k_a$ | TMZ absorption rate | $\frac{1}{day}$ | 139.2 | Fixed[7] |
| $CL$ | TMZ clearance rate | $\frac{L}{day}$ | 240 | Fixed[7] |
| $V_D$ | TMZ volume of distribution in the central compartment | L | 30.3 | Fixed[7] |
| $k_{23}$ | TMZ plasma to CSF transfer rate | $\frac{1}{day}$ | 0.01728 | Fixed[7] |
| $k_{32}$ | TMZ CSF to plasma transfer rate | $\frac{1}{day}$ | 18.24 | Fixed[7] |
| $V_p$ | TMZ volume of distribution in the CSF | mL | 140 | Fixed[7] |
| $E_{max}$ | TMZ maximal effect | $\frac{1}{day}$ | 0.30 | Fit[44] |
| $h$ | TMZ Hill coefficient | Unit-less | 0.5682 | Fit[44] |
| $IC50_{TMZ}$ | TMZ IC50 | $\frac{mg}{ml}$ | 0.0011 | Fit[44] |

**Supplementary Table 9. Parameters of the chemotherapy treatment.**

| Parameter | Definition | Units | Estimated value | Reference |
|---|---|---|---|---|
| $\mu_I$ | Binding rate of anti-PD-1 to PD-1 receptors | $\frac{\text{ml}}{\text{pg} \cdot \text{day}}$ | $3.8186 \cdot 10^{-6}$ | Calculated[13] |
| $\mu_P$ | Anti-PD-1 blocking rate of PD-1 receptors | $\frac{\text{ml}}{\text{mg} \cdot \text{day}}$ | $3.8186 \cdot 10^{3}$ | Calculated[13] |
| $\delta_I$ | Natural clearance of anti-PD-1 | $\frac{1}{\text{day}}$ | 0.0257 | Calculated[37] |

**Supplementary Table 10. Parameters of the anti-PD-1 immunotherapy treatment.**

### Initial conditions

To model cancer initiation, we assumed an initial tumour volume of 5 cubic millimeters meaning that, at $t = 0$, $G(0) = 5 \cdot 10^6$ cancer cells. Initial cell-to-cell ratios were obtained from Surendran et al.[27] and Karimi et al.[49] who analysed glioblastoma biopsies and resections using imaging mass cytometry. Accordingly, we assumed that TAMs were mostly inactivated at cancer initiation and hence that there is initially 1 anti-tumoural TAM per $10^6$ resident TAMs. With an average M2:M1 ratio of 3.2 in glioblastomas[34], this translates to an initial 3.2 M2 count. From this value, we calculated the fraction of resident TAMs that acquire a pro-tumoral phenotype upon activation $(q)$. We assumed the initial conditions in our model to be the conditions at the no-tumour equilibrium, which allowed us to calculate the constant source of resident TAMs in the TME $(\lambda_M)$ and the maximal activation rate of CD8+ T cells by the innate immune system $(\eta_1)$. The initial concentration of unbound PD-1 receptors was calculated according to

$$PD1_U(0) = \rho_T \cdot T(0), \tag{S28}$$

with the initial concentration of PD-L1 receptors is given by

$$PDL1(0) = \rho_G \cdot G(0) + \rho_M \cdot \big(M(0) + M_1(0) + M_2(0)\big), \tag{S29}$$

Initial conditions are listed in **Supplementary Table 11**.

| Initializations | Definition | Value | Units |
|---|---|---|---|
| $G(0)$ | Initial glioblastoma cell population | 5 | $10^6$cells |
| $T(0)$ | Initial CD8+ T cell population | 0.0320 | $10^6$cells |
| $M(0)$ | Initial resident TAM population | 1.2516 | $10^6$cells |
| $M_1(0)$ | Initial anti-tumoral TAM $(M_1)$ population | $1.2516 \cdot 10^{-6}$ | $10^6$cells |
| $M_2(0)$ | Initial pro-tumoral TAM $(M_2)$ population | $4.0051 \cdot 10^{-6}$ | $10^6$cells |
| $PD1_U(0)$ | Initial anti-PD-1 unbound PD-1 receptor concentration in the TME | 0.0416 | pg/ml |
| $PDL1(0)$ | Initial PD-L1 concentration in the TME | 170.3663 | pg/ml |
| $Dd(0)$ | Initial dead and damaged glioblastoma cell population | 0 | $10^6$cells |

**Supplementary Table 11. Initial conditions of cellular populations considered in the model.**

### Tumour growth in M1- and M2-knockout models

To further analyse the effect of TAM polarisation on tumour growth, we simulated M1- and M2-knockouts in two contexts: 1) during treatment and 2) from cancer initiation. Knockouts were simulated by setting the given population to 0 during the period of interest. To ensure no cells acquired the knocked-out phenotype, we set $q = 1$ and $\mu_{M2} = 0$ during M1-knockout, and

$q = 0$ and $\mu_{M1} = 0$ during M2-knockout. With a high M2:M1 ratio on average, glioblastoma growth in our original SOC+ICI model resembles that of the complete M2-knockout model (**Supplementary Figure 3A**), even though the presence of a few antitumoral TAMs results in more CD8+ T cell cytotoxic activity (**Supplementary Figure 3C**). Since CD8+ T cell activation by the innate immune system is dependent on the M2:M1 ratio, the complete M1-knockout scenario (i.e., from cancer initiation) resulted in no CD8+ T cell activation by TAMs (**Supplementary Figure 3B**), which translates to low levels of CD8+ T cells and very few tumour cells killed by the adaptive immune system (**Supplementary Figure 3C**, inset). In the complete M2-knockout model, most TAMs are antitumoral which results in high CD8+ T cell activation by the innate immune system (**Supplementary Figure 3B**). Accordingly, our model predicted high levels of CD8+ T cell cytotoxicity (**Supplementary Figure 3C**) and tumour stabilization below the lethal threshold (**Supplementary Figure 3A**).

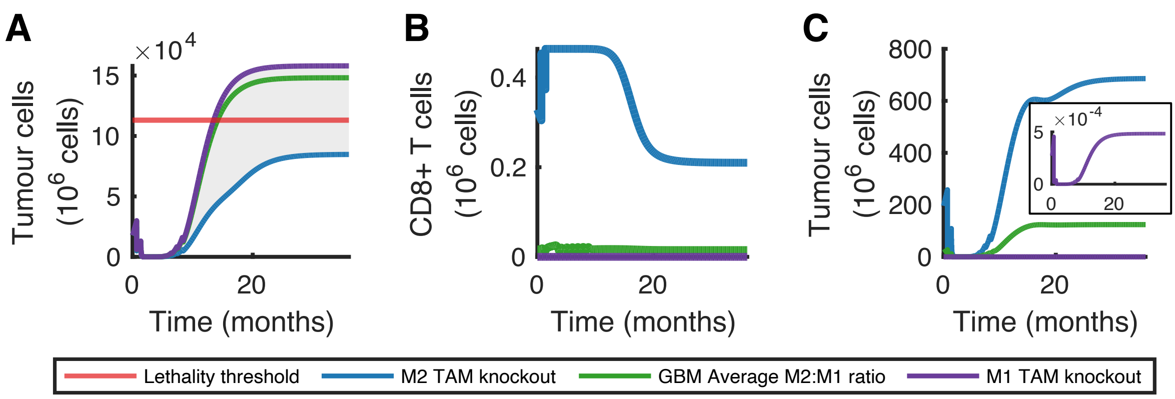


**Supplementary Figure 3. Glioblastoma growth under complete M1- or M2- knockout during combination SOC+ICI treatment. A)** In the complete M1-knockout model (purple), the tumour grows rapidly, reaching the lethal threshold (red) after 13.51 months under SOC+ICI treatment. On the contrary, the tumour was predicted to stabilize below the lethal threshold under complete M2-knockout (blue). Since a glioblastoma has a high M2:M1 ratio on average, its growth (green) is similar to that in the M2-knockout model. **B)** Number of CD8+ T cells activated by TAMs over time during SOC+ICI treatment. **C)** Number of glioblastoma cells killed by CD8+ T cells over time during SOC+ICI treatment. **A-C)** Time refers to time after diagnosis.

### Parameter sensitivity analysis

To identify the parameters with the most impact on model predictions, we performed a global sensitivity analysis (GSA) using Extended Fourier Amplitude Sensitivity Test (eFAST). eFAST assigns to each parameter $p$ a first-order sensitivity index $\left(S_p\right)$ and and total-order sensitivity index $\left(S_{T_p}\right)$. The first-order sensitivity index represents the fraction of model output variance explained by the variation of $p$[50]. The total-order sensitivity index accounts for higher-order, nonlinear interactions between $p$ and the other other parameters[50] and is defined as the remaining variance after we remove the first-order sensitivity index of the other parameters. We evaluated the sensitivity of our model to the following outputs: 1) intrinsic growth of the tumour ($\beta$), 2) activity of the adaptive ($\alpha_{AT}$ and $\alpha_2$) and 3) innate $\left(\eta_1, \mu_{M_1}, \mu_{M_2}, q \text{ and } \lambda_M\right)$ immune systems, and 4) ICI binding efficacy ($\mu_P$), see **Supplementary Figure 4**. To reduce the dimensionality of the parameter set, we only varied the maximal rate and not the half-effect concentration in interaction terms defined in Michaelis-Menten relationships as we assumed that if our model was sensitive to the maximal rate, then it was also sensitive to its corresponding interaction. Following Marino et al.[50], we used 5 search curves with 1,000 samples per search curve, at most four Fourier coefficients, and uniform distributions ranging from 0.5-1.5-times the baseline value of each parameters of interest, as we had no information on predefined distributions. For the parameter $q$, we used a uniform distribution ranging from 0 to 1.

We measured changes to model outputs by considering the tumour burden after SOC and the treatment supplementary efficacy conferred by the addition of nivolumab to the SOC, i.e., $\mathrm{S}(\mathrm{SOC}+\mathrm{ICI})$. To identify statistically significantly sensitive parameters, we performed a two-sample t-test using a dummy variable that does not appear in our model[50]. We defined sensitive

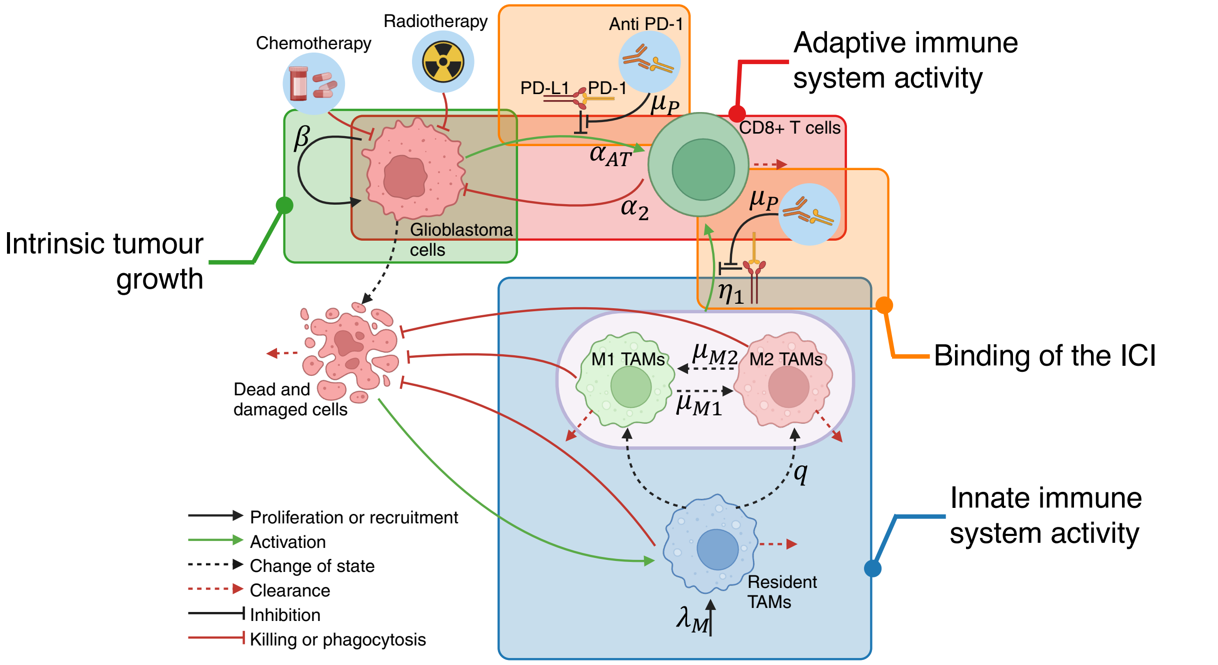


**Supplementary Figure 4. Global sensitivity analysis paradigm.** Schematic of the model highlighting the interactions of interest for the global sensitivity analysis (GSA). We evaluated the sensitivity of our model to the 1) intrinsic tumour growth rate ($\boldsymbol{\beta}$), 2) antigenicity of tumour cells ($\boldsymbol{\alpha_{AT}}$), 3) cytotoxic activity of CD8+ T cells ($\boldsymbol{\alpha_2}$), 4) activation of CD8+ T cells by activated TAMs ($\boldsymbol{\eta_1}$), 5) phenotype switching rates between $\boldsymbol{M_1}$ and $\boldsymbol{M_2}$ TAMs $\left(\boldsymbol{\mu_{M_1}}\ \textbf{\textit{and}}\ \boldsymbol{\mu_{M_2}}\right)$, 6) TME bias towards M2-activated TAMs ($\boldsymbol{q}$), 7) constant source of resident TAMs in the TME ($\boldsymbol{\lambda_M}$), and 8) ICI binding efficacy ($\boldsymbol{\mu_P}$). Made with BioRender.

parameters as those with $S_{T_p} > S_{T_{\mathrm{dummy}}}$, and with their $S_{T_p}$ distributions due to the resampling significantly different from that of the dummy parameter, at the $\alpha = 1\%$ significance level[50].

A complete list of p-values from the global sensitivity analysis are reported in **Supplementary Table 12**. Both the tumour burden after SOC and nivolumab efficacy $\left(S(\mathrm{SOC}+\mathrm{ICI})\right)$ were found to be significantly sensitive to the intrinsic growth of the tumour cells, as well as to the activity of the adaptive and innate immune systems. However, $S(\mathrm{SOC}+\mathrm{ICI})$ was not found to be sensitive to the ICI blocking efficacy ($\mu_P$), further suggesting that clinical failures are not due to the mechanism of action of the ICI but rather to interactions within the TME. This finding is consistent with other studies that attribute these failures to a lack of recruitment of CD8+ T cells[27,51] and is coherent with our result that shows an increase in survival time when we introduce a constant source of CD8+ T cell recruitment during treatment (**Figure 3B** in the Main Text). In absence of a treatment inhibiting PD-1/PD-L1 suppression of CD8+ T cells, the tumour burden after SOC was not determined to be significantly sensitive to the antigenicity of tumour cells ($\alpha_{AT}$), but it was found to be sensitive to the rate of CD8+ T cell-induced glioblastoma rate (**Supplementary Figure 5A**).

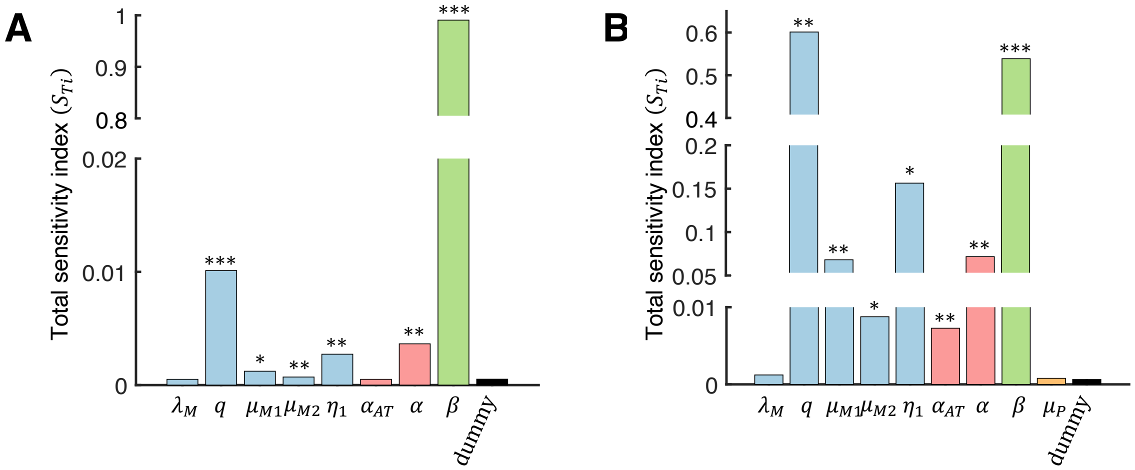


**Supplementary Figure 5. Global sensitivity analysis results. A)** Tumour burden after SOC. **B)** Supplementary efficacy conferred by the addition of nivolumab to SOC (i.e., S(SOC+ICI)) was found to be statistically significantly sensitive to the tumour's intrinsic growth rate (**β**, green), and the activity of the innate (light blue) and adaptive (light red) immune systems. *: p-value $< \mathbf{10^{-2}}$, **: p-value $< \mathbf{10^{-4}}$, ***: p-value $< \mathbf{10^{-8}}$

| **p-value (*t*-test)** | $\lambda_M$ | $q$ | $\mu_{M1}$ | $\mu_{M2}$ | $\eta_1$ | $\alpha_{AT}$ | $\alpha_2$ | $\beta$ | $\mu_P$ |
|---|---|---|---|---|---|---|---|---|---|
| **Tumour burden after SOC** | 0.7717 | $1.3963e-09$ | 0.0006 | $2.3595e-06$ | $1.0689e-05$ | 0.7506 | $4.8063e-05$ | $1.6356e-16$ | NA |
| **SE(SOC+ICI)** | 0.0187 | $1.8919 \times 10^{-7}$ | $5.9302 \times 10^{-7}$ | 0.0048 | 0.0003 | $3.5042 \times 10^{-5}$ | $1.4514 \times 10^{-5}$ | $8.4051 \times 10^{-9}$ | 0.1700 |

**Supplementary Table 12. Results of the global sensitivity analysis.** To identify the sensitivity of the tumour burden after SOC and nivolumab's supplementary efficacy (i.e., $\boldsymbol{S(SOC+ICI)}$) to model parameters, we performed a *t*-test comparing each parameter's total sensitivity index ($\boldsymbol{S_{Ti}}$) distribution with that of a dummy variable. The output was considered sensitive to a parameter if the parameter's $\boldsymbol{S_{Ti}}$ was larger than the dummy's and if its distribution was significantly different from that of the dummy at the $\boldsymbol{\alpha = 1\%}$ significance level. NA: Not applicable, as ICI binding is not included in SOC.

### Tumour burden after treatment when reducing TAM numbers in TME

To investigate the potential of TAM-targeting treatments reducing TAMs in the TME, we first mimicked the effect of an anti-CSF1R monoclonal antibody (mAB) by simulating a treatment that increased the rate of TAM death. Compared to SOC, this strategy in combination with the SOC failed to reduce the tumour burden and even led to a small increase in tumour cell count after treatment (**Supplementary Figure 6A**). On the contrary, increasing only the M2 TAM death rate resulted in a post-treatment tumour cell count decrease of over 12% (**Supplementary Figure 6C**). This suggests that failures of anti-CSF1R mABs such as Emactuzumab could be attributed to a lack of specificity, as they do not specifically target pro-tumoral TAMs[52].

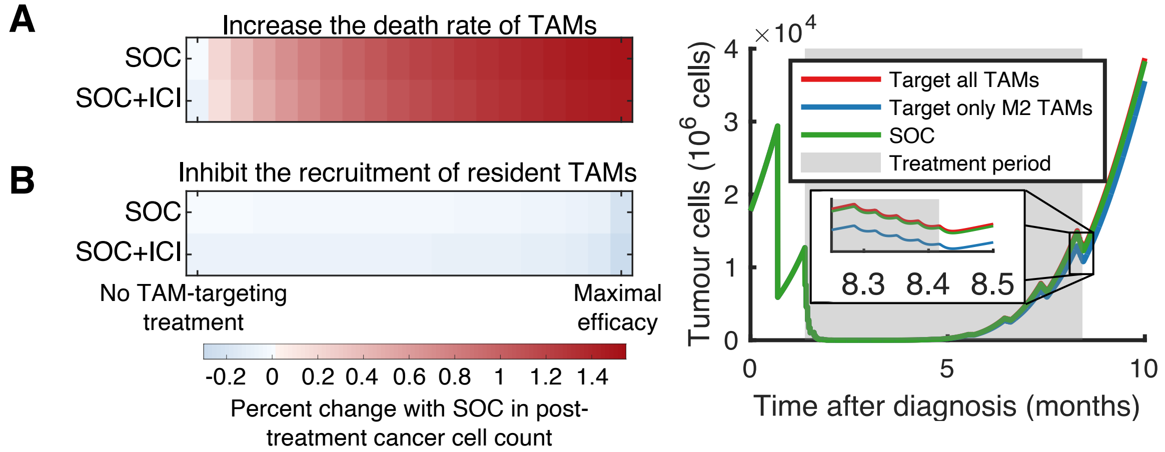


**Supplementary Figure 6. Reducing TAMs in TME fails to reduce tumour burden after treatment. A-B)** Efficacy of indicated TAM-targeting treatment in combination with SOC (top row) and SOC+ICI (bottom row), compared to SOC. We varied the efficacy of the TAM-targeting agent between no effect and maximal, as detailed in SI section **Targeting tumour-associated macrophages and microglia**. Heatmaps show the percent change in tumour cell count one week after treatment compared to SOC. Increasing the death rate of all TAMs (A) resulted in a small increase in tumour cell count after treatment, while inhibiting the recruitment of resident TAMs in the tumour micro-environment (B) did not significantly reduce post-treatment tumour burden. Both strategies were not found to increase the activation of CD8+ T cells by the innate immune system. **C)** Increasing only the death rate of M2 TAMs decreased the M2:M1 ratio in the TME. This added specificity increased the activation of CD8+ T cells by the innate immune system and resulted in a post-treatment tumour burden decrease of over 12%.

Reducing the number of TAMs can also be achieved by inhibiting the recruitment of resident TAMs in the TME with an antibody that binds to CCL2, for example. We simulated a similar treatment by decreasing the constant source of resident TAMs in our model. The M2:M1 ratio was not impacted in this scenario and, as such, this strategy only results in a minimal decrease of tumour burden after treatment, even with maximal efficacy (**Supplementary Figure 6B**). This is coherent with the results of our GSA that showed that both tumour burden after SOC and nivolumab's efficacy (i.e., $S(\text{SOC} + \text{ICI})$) are not sensitive to the constant source of resident TAMs in our model ($\lambda_M$), see **Supplementary Figure 5**.

### Virtual individual parameters

To generate virtual patients, we first defined log-normal distributions for each of the parameters varied in our virtual population. In a normal distribution, 99.7% of the data points will be between 3 standard deviations of the mean. For each parameter $p$ varied in the VP cohort, we set its mean value $\hat{\mu}_p$ to its baseline value, then defined $p_1$ and $p_2$ as the limits within which 99.7% of the values should lie. In this way, the natural logarithms of $p$ followed a normal distribution with parameters $\mu_p$ and $\sigma_p$ given by:

$$\mu_p = \log \hat{\mu}_p, \tag{S30}$$

$$\sigma_p = \min\left(\frac{|\mu_p - \log(p_1)|}{3}, \frac{|\mu_p - \log(p_2)|}{3}\right). \tag{S31}$$

We also defined intervals of definition for every parameter varied to exclude unrealistic samples.

Our GSA showed that the tumour cell count after SOC with or without nivolumab was statistically significantly sensitive to the intrinsic growth of the tumour cells ($\beta$), as well as to the activity of the adaptive and innate immune systems (**Supplementary Figure 6**). Since $\beta$ is a growth rate, it must be positive and is defined in the interval $[0, +\infty)$ (**Supplementary Table 13**). Thus, we varied the intrinsic growth rate of the tumour cells $(\beta)$ by finding $\beta_1$ $(\beta_2)$ such that the simulated survival time without treatment decreased (increased) by one month. We found that $\beta_1 = 0.0157$ (i.e., -0.9865 months) and $\beta_2 = 0.0111$ (i.e., +1.0225 month).

Both the antigenicity of tumour cells $(\alpha_{AT})$ and the killing rate of the tumour cells by the CD8+ T cells $(\alpha_2)$ in the adaptive response had total sensitivity indices statistically significantly different from that of the dummy parameter for $S(\text{SOC} + \text{ICI})$, with p-values within the same order of magnitude. Surendran et al. showed that higher levels of CD8+ T cell in TME lead to better ICI responses[27] Similarly, our model predicted that increasing CD8+ T cell recruitment during treatment increases survival (**Figure 1B** in the Main Text). Thus, we varied $\alpha_{AT}$ and kept $\alpha_2$ fixed. $\alpha_{AT}$ must be positive and is defined in the interval $[0, +\infty)$ (**Supplementary Table 13**). Varying $\alpha_{AT}$ one order of magnitude under and over its baseline value was predicted to lead to survival times between 14.0533 and 16.1198 months under SOC, which is coherent with clinical observations[1].

Finally, for the innate immune system, the parameter $q$ (i.e., the TME bias towards a pro-tumoral TAM phenotype) was the parameter found to have the largest total sensitivity index (see **Supplementary Figure 6**). Since $q$ is directly linked to the M2:M1 ratio (see **Parameter estimations** section above), varying it to generate our virtual population allowed us to study the importance of the M2:M1 ratio as a prognostic factor. An average M2:M1 ratio of 3.2 in glioblastoma patients[34] translates to a $q$ value of 0.7619. To define a distribution for $q$, we assumed that the percentage of TAMs with the M2 phenotype was between 5% (i.e. M2:M1=1:19) and 95% (i.e. M2:M1=19:1) (**Supplementary Table 13**). Resulting distributions are shown in **Supplementary Figure 8**.

| Parameter $p$ | $\widehat{\mu_p}$ | $p_1$ | $p_2$ | Interval of definition |
|---|---|---|---|---|
| $\beta$ | 0.013 | 0.0111 | 0.0157 | $[0, +\infty)$ |
| $\alpha_{AT}$ | 0.0375 | 0.00375 | 0.375 | $[0, +\infty)$ |
| $q$ | 0.7619 | 0.05 | 0.95 | $[0,1]$ |

**Supplementary Table 13. Definitions of normal distributions for the natural logarithms of the intrinsic growth rate of tumour cells $(\boldsymbol{\beta})$, the antigenicity of tumour cells $(\boldsymbol{\alpha_{AT}})$ and the TME bias towards M2-activated TAMs $(\boldsymbol{q})$.** For every parameter $\boldsymbol{p}$ varied in the population, $\boldsymbol{log(p)} \sim \boldsymbol{\mathcal{N}(\mu_p, \sigma_p)}$ with $\boldsymbol{\mu_p = log(\widehat{\mu}_p)}$ and $\boldsymbol{\sigma_p = min\left(\frac{|\mu_p - log(p_1)|}{3}, \frac{|\mu_p - log(p_2)|}{3}\right)}$. When sampling a parameter $\boldsymbol{p_i}$ to include it in the set corresponding to the $\boldsymbol{i^{th}}$ virtual patient, we sampled the natural logarithm $\boldsymbol{log(p_i)}$ from $\boldsymbol{\mathcal{N}(\mu_p, \sigma_p)}$ and then calculated $\boldsymbol{p_i = exp(log\, p_i)}$. To be included in the set of parameters, $\boldsymbol{p_i}$ must be contained in the interval of definition for $\boldsymbol{p}$.

The population pharmacokinetic model established by Ostermann et al.[7] defined log-normal distributions for the TMZ clearance rate $CL$, its absorption rate $k_a$, and its plasma-to-CSF transfer rate $k_{23}$ (**Supplementary Table 14**). We integrated these distributions in our virtual population to simulate varying TMZ pharmacokinetics in the population.

| **Parameter** | **Definition** | **Units** | **Population mean** | **Inter-individual variability (IIV)** | **Standard deviation** |
|---|---|---|---|---|---|
| $k_{CL}$ | TMZ clearance | L/h | 10 | 4.7% | 0.0470 |
| $k_a$ | TMZ absorption rate | 1/h | 5.8 | 111% | 0.1649 |
| $k_{23}$ | TMZ plasma-to-CSF transfer rate | 1/h | $7.2 \cdot 10^{-4}$ | 16.6% | 0.8961 |

**Supplementary Table 14. Log-normal distributions for the pharmacokinetics of TMZ, defined by the Pop-PK model established by Ostermann et al.[7]** The standard deviation was calculated as $\boldsymbol{sd} = \sqrt{\boldsymbol{log}(\text{IIV}^2 + \mathbf{1})}$.

We then sampled 900 virtual patients from the distributions for $\beta$, $\alpha_{AT}$, $q$, $k_{CL}$, $k_a$ and $k_{23}$ (**Supplementary Figure 7**).

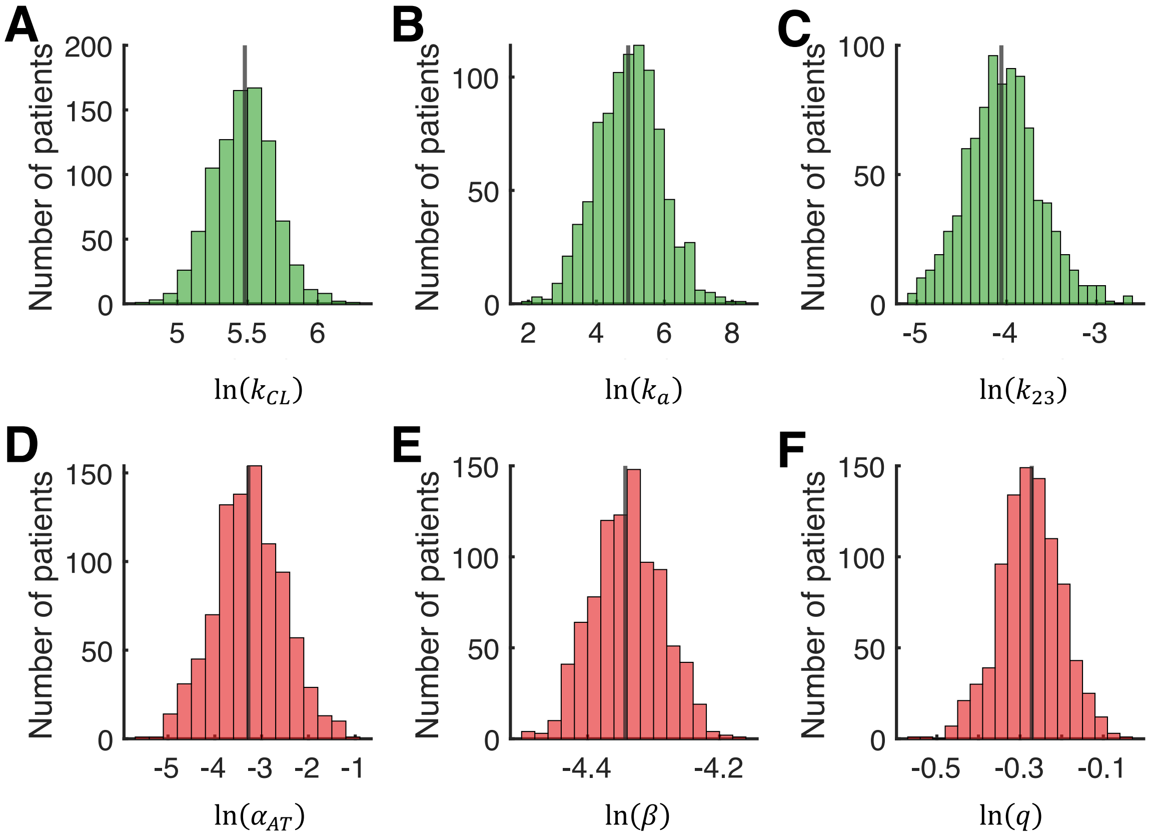


**Supplementary Figure 7. Virtual patient distributions.** Parameters were sampled from log-normal distributions. The antigenicity of tumour cells ($\boldsymbol{\alpha_{AT}}$), the intrinsic tumoral growth rate ($\boldsymbol{\beta}$) and the TME bias towards a pro-tumoral TAM phenotype ($\boldsymbol{q}$) were chosen to be varied in the virtual population based on results from our global sensitivity analysis. The TME clearance rate ($\boldsymbol{k_{CL}}$), absorption rate ($\boldsymbol{k_a}$) and plasma-to-CSF transfer rate distributions come from a population pharmacokinetic model by Ostermann et al.[7]. Solid black lines: natural logarithm of the parameter value chosen as the mean in the virtual population.

### Virtual clinical trial of SOC combined with nivolumab

We simulated SOC with or without the anti-PD-1 ICI nivolumab in our virtual patient cohort (see Section **Virtual individual parameters** and **Figure 2A** in Main Text). Our model predicted that with SOC, tumours would reach the lethal threshold between 12.4085 and 16.2798 months after diagnosis, and between 12.4421 and 18.0295 months with SOC+ICI (**Supplementary Figure 8B**). Given that nivolumab has not been shown to significantly improve glioblastoma overall survival[53–55], our model successfully captures these past clinical trial failures. To understand the role of each parameter used to generate the virtual population played on survival, we compared Kaplan-Meier curves between the virtual patients with a values smaller than the median and those with a values larger than the median for a given parameters (**Supplementary Figure 8C-H**). For the TMZ pharmacokinetics, survival was significantly higher in the low clearance rate ($k_{CL}$) group than in the high $k_{CL}$ group ($p < 10^{-11}$, log-rank test), and in the high plasma-to-CSF transfer rate ($k_{23}$) group than in the low $k_{23}$ group ($p < 10^{-6}$, log-rank test). This result is physiologically coherent as a lower clearance rate results in more drug exposure, and a higher plasma-to-CSF transfer rate translates in more drug reaching the site of action (i.e. CSF). A higher intrinsic tumour growth rate ($\beta$) was also a predictor of poor outcome, with virtual patients with $\beta$ values larger or equal to the median predicted to have the worst prognoses. This underlies the role on survival played by the aggressive growth of this tumour and emphasizes the need to establish effective treatment strategies with a direct effect on this growth.

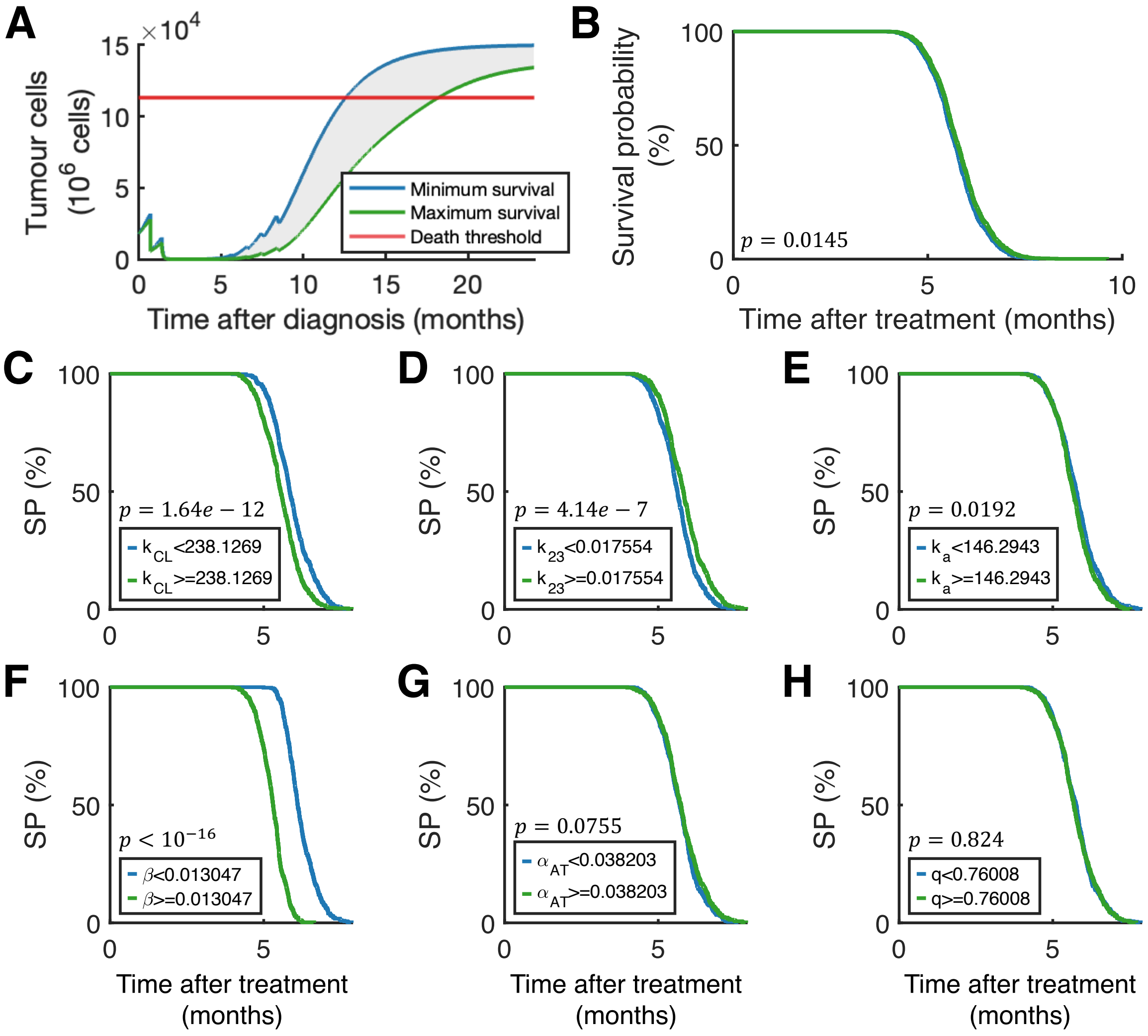


**Supplementary Figure 8. Virtual clinical trial of SOC combined with nivolumab. A)** Tumours in the virtual population reach the lethality threshold between 12.4421 (blue) and 16.2798 (green) months after diagnosis. **B)** Nivolumab was not predicted to significantly increase survival (p-value = $\mathbf{0.0145}$, log-rank test). **C-H)** Kaplan-Meier curves showing the survival probability (SP) with SOC based on the six parameters varied in the virtual cohort. Each curve compares those virtual with values below (blue) and those with values larger or equal (green) to the median value in the log-normal distribution. The TMZ clearance rate $\boldsymbol{k_{CL}}$ (C), the TMZ plasma-to-CSF transfer rate $\boldsymbol{k_{23}}$ (D) and the intrinsic tumoral growth rate $\boldsymbol{\beta}$ (F) separate virtual patients into two groups with significantly different survival curves at the $\boldsymbol{\alpha} = \mathbf{0.01}$ level. Survival is not significantly different virtual patients with low or high TMZ absorption rate $\boldsymbol{k_a}$ (E), antigenicity of tumour cells $\boldsymbol{\alpha_{AT}}$ (G) or TME bias towards M2 TAM phenotype $\boldsymbol{q}$ (H).

#### Statistical analysis of VCT to enhance M1 TAM phagocytic activity

We simulated the enhancement of M1 TAM phagocytic activity in our 900 virtual patients; predicted survival times for each virtual patient were used to generate Kaplan-Meier curves. We used the survival analysis function *MatSurv*[56] in MatlabR2024a[46] to calculate the hazard ratio (HR) between two survival curves and to perform log-rank tests to identify significantly different survival curves. Survival was found to be significantly higher in treatment combinations that included the enhancement of M1 TAM phagocytic activity (see **Supplementary Table 15**).

| SOC+TAMT vs SOC | SOC+TAMT vs SOC+ICI | SOC+TAMT+ICI vs SOC | SOC+TAMT+ICI vs SOC+ICI |
|---|---|---|---|
| $0.227\ (0.201 - 0.255)$ $p - \text{value} < 10^{-16}$ | $0.247\ (0.22 - 0.278)$ $p - \text{value} < 10^{-16}$ | $0.221\ (0.196 - 0.249)$ $p - \text{value} < 10^{-16}$ | $0.239\ (0.212 - 0.268)$ $p - \text{value} < 10^{-16}$ |

**Supplementary Table 15. Hazard ratio (95% confidence interval) between survival curves of treatment combinations with and without enhancement of phagocytosis of glioblastoma cells by M1 TAMs.** In all cases, the combination treatment including the TAM-targeting treatment (TAMT) were predicted to significantly improve the survival probability and median overall survival. $\boldsymbol{p}$-values smaller than numerical precision ($\mathbf{10^{-16}}$) were reported as $\boldsymbol{p} - \textbf{value} < \mathbf{10^{-16}}$.

To identify patient-specific characteristics distinguishing best from worst responders to a treatment restoring M1 phagocytic activity in combination with SOC, we normalized each patient's supplementary efficacy by their untreated tumour growth over the same period (see **Statistical Analysis** in the Main Text). We then segregated the virtual patient cohort into quartiles based on their normalized $S$ values and defined the worst and best responders as the patients in the first and third quartiles, respectively. We also used the overlapping index $\hat{\eta}$ as defined by Pastore and Calcagni[57] as an effect size measure. We identified parameters distinguishing best from worst responders if a two-sided Wilcoxon rank sum test was statistically significant with $\hat{\eta} < 0.5$ (see **Statistical Analysis** in Main Text). While the intrinsic tumour growth rate $(\beta)$, antigenicity of tumour cells $(\alpha_{AT})$ and TME bias towards M2 activated TAMs $(q)$ all have $p$-values smaller than 0.05, only $q$ has an overlapping index smaller than 0.5 and, thus, distinguish best from worst responders (see **Supplementary Table 16**).

| Treatment | Parameter | $\boldsymbol{p}$-value (Wilcoxon rank sum test) | Overlapping index $\hat{\boldsymbol{\eta}}$ |
|---|---|---|---|
| SOC+TAMT3 | $CL$ | 0.0538 | 0.0273 |
| | $k_{23}$ | 0.1329 | 0.937 |
| | $k_a$ | 0.5507 | 0.915 |
| | $\beta$ | $7.8436e - 07$ | 0.808 |
| | $\alpha_{AT}$ | 0.0005 | 0.793 |
| | $q$ | $< 10^{-16}$ | 0.0273 |

**Supplementary Table 16. $\boldsymbol{p}$-values and overlapping indices $\hat{\boldsymbol{\eta}}$ between the best and worst responders.** Distributions for $\boldsymbol{\beta}$, $\boldsymbol{\alpha_{AT}}$ and $\boldsymbol{q}$ were all found to be statistically significantly different between the worst and best responders, but only the TME bias towards the pro-tumoral TAM phenotype ($\boldsymbol{q}$) had an overlapping index smaller than $\mathbf{0.5}$ (shaded cells). Thus, we defined it as distinguishing best from worst responders. $\boldsymbol{p}$-values smaller than numerical precision are noted as $<\mathbf{10^{-16}}$.

### Parameter distribution of M1-biased and M2-biased virtual populations

To generate M1- and M2-biased virtual populations, we set the distributions of $k_{CL}$, $k_a$, $k_{23}$, $\alpha_{AT}$, and $\beta$ as in our original 900 virtual cohort and defined a new log-normal distribution for $q$, with either a mean value of $q = 0.25$ and $q = 0.75$, respectively. To ensure that 99.7% of our virtual populations would have $q$ values within the intervals $[0{,}0.5]$ (M1-biased) and $[0.5{,}1]$ (M2-biased), we calculated the standard deviation of the natural logarithm of $q$ using with **Eqs. (S30)** and **(S31)**, with $\widehat{\mu_p} = 0.25$, $p_1 = 0$ and $p_2 = 0.5$ for the M1-biased virtual cohort, and $\widehat{\mu_p} = 0.75$, $p_1 = 0.5$ and $p_2 = 1$ for the M2-biased virtual cohort. 450 VPs were sampled from the parameter distributions for each of the virtual cohorts (**Supplementary Figure 9** and **Supplementary Figure 10**).

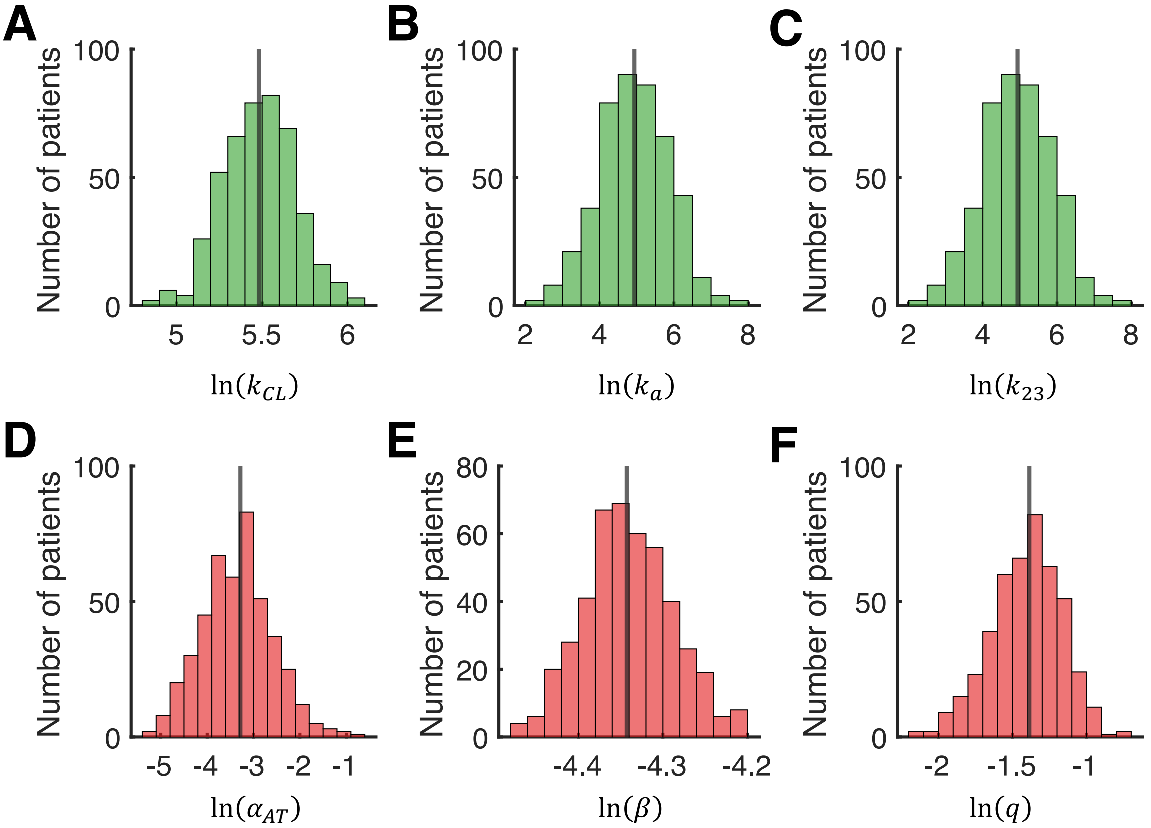


**Supplementary Figure 9. Parameter distributions for the M1-biased virtual population.** Parameters were sampled from log-normal distributions to generate 450 virtual patients with a TME biased towards a M1 TAM phenotype. The TME clearance rate $\boldsymbol{k_{CL}}$ (A), absorption rate $\boldsymbol{k_a}$ (B) and plasma-to-CSF transfer rate $\boldsymbol{k_{23}}$ (C) distributions come from a population pharmacokinetic model by Ostermann et al.[7]. The antigenicity of tumour cells $\boldsymbol{\alpha_{AT}}$ (D) and the intrinsic tumoral growth rate $\boldsymbol{\beta}$ (E) distributions were defined as in our virtual general population (see **Virtual individual parameters**). We follow the same methodology to generate a distribution for the TME bias towards the M2 TAM phenotype $\boldsymbol{q}$ (F), but with $\boldsymbol{\widehat{\mu_q} = 0.25}$, $\boldsymbol{q_1 = 0}$ and $\boldsymbol{q_2 = 0.5}$ to model a M1-biased virtual population. Solid black lines: natural logarithm of the parameter value chosen as the mean in the virtual population.

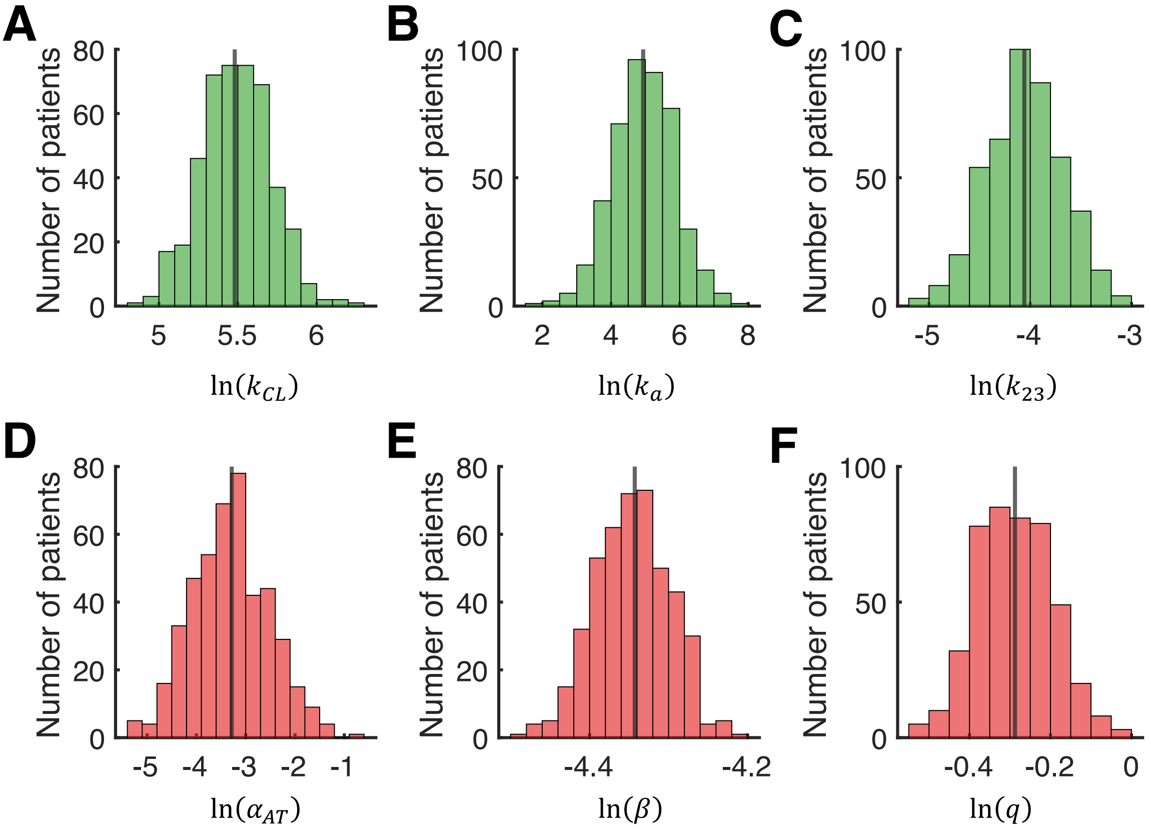


**Supplementary Figure 10. Parameter distributions for the M2-biased virtual population.** Parameters were sampled from log-normal distributions to generate 450 virtual patients with a TME biased towards a M2 TAM phenotype. The TME clearance rate $\boldsymbol{k_{CL}}$ (A), absorption rate $\boldsymbol{k_a}$ (B) and plasma-to-CSF transfer rate $\boldsymbol{k_{23}}$ (C) distributions come from a population pharmacokinetic model by Ostermann et al.[7]. The antigenicity of tumour cells $\boldsymbol{\alpha_{AT}}$ (D) and the intrinsic tumoral growth rate $\boldsymbol{\beta}$ (E) distributions were defined as in our virtual general population (see **Virtual individual parameters**). We follow the same methodology to generate a distribution for the TME bias towards the M2 TAM phenotype $\boldsymbol{q}$ (F), but with $\widehat{\boldsymbol{\mu_q}} = \mathbf{0.75}$, $\boldsymbol{q_1} = \mathbf{0.5}$ and $\boldsymbol{q_2} = \mathbf{1}$ to model a M2-biased virtual population. Solid black lines: natural logarithm of the parameter value chosen as the mean in the virtual population.
